# Variants at the ADAMTS13, BGALT5, SSBP2 and TKT Loci are associated with Post-term birth

**DOI:** 10.1101/153833

**Authors:** William Schierding, Jisha Antony, Ville Karhunen, Marja Vääräsmäki, Steve Franks, Paul Elliott, Eero Kajantie, Sylvain Sebert, Alex Blakemore, Julia A. Horsfield, Marjo-Riitta Järvelin, Justin M. O’Sullivan, Wayne S. Cutfield

## Abstract

Gestation is a crucial timepoint in human development. Deviation from a term gestational age correlates with both acute and long-term adverse health effects for the child. Both being born pre and post-term, *i.e.* having short and long gestational ages, are heritable and influenced by the pre- and perinatal environment. Despite the obvious heritable component, specific genetic influences underlying differences in gestational age are poorly understood. Here we identify one globally significant intronic genetic variant within the *ADAMTS13* gene that is associated with prolonged gestation in 9,141 white European individuals from the 1966 and 1986 Northern Finland birth cohorts. Additional variants that reached suggestive levels of significance were identified within introns at the *TKT,* and *ARGHAP42* genes, and in the upstream (5’) intergenic regions of the *B3GALT5* and *SSBP2* genes. The variants near the *ADAMTS13, B3GALT5, SSBP2* and *TKT* loci are linked to alterations in gene expression levels (cis-eQTLs). Luciferase assays confirmed the allele specific enhancer activity for the *BGALT5* and *TKT* loci. Our findings provide the first evidence of a specific genetic influence associated with prolonged gestation.

## Introduction

Gestation is a crucial period of human development. Being born too early (preterm, < 37 weeks gestation) or too late (post-term, ≥42 weeks gestation) can have significant acute and long-term health consequences [1]. While preterm birth has received substantial attention[2–4], post-term birth has been scantily explored despite approximately 3-5% of all births each year being post-term[5]. Prolonged gestation poses a unique set of acute and long-term adverse health outcomes, including an increased need for intervention during labour and risk factors for truncal obesity, insulin resistance, altered lipids and elevated blood pressure [6, 7]. Thus, there is a vital need to understand the role of genetic variants on post-term birth.

The acute health risks, for both the mother and the child, of being born post-term are well documented (*for review see* [8]). Consequently, induction of birth at or before 41 weeks gestation is recommended in order to reduce the acute risks associated with post-term birth[5, 8]. As a result, approximately 25% of the routine inductions in Australia in 2010 were primarily performed to prevent prolonged pregnancy. However, the rules regarding the decision to induce labour are not consistently applied across different hospitals, reflecting the influence of opinions of individual practitioners and differing staff routines[9]. Despite this, induction remains an excellent intervention, and its application has reduced post-term births from approximately 20% of all births in the 1960’s[10] to the modern-day rate of under 5%[5, 11]. However, there is a possibility that the long-term risks associated with post-term birth are a part of the genetically-informed trajectory for induced individuals. In this case, induction would not change this aspect of the biology of post-term individuals. Family and twin studies attribute 25-40% of the variation in gestational age to genetic factors [12–19] with fetal (26%) and maternal (21%) factors each explaining nearly half of this variation [18]. Thus, there is a large population of individuals who have “post-term potential”[20] and possibly face the long-term health risks of post-term birth without actually having been born post-term (due to pregnancy ending from obstetric management).

Evidence supports the hypothesis that it is the fetus that determines the timing of labor rather than the mother. Therefore, we have investigated the heritability and the genetic architecture of gestational age in 9,141 Northern Finnish (white European) individuals (1,167 post-term) across two birth cohorts (Northern Finland Birth Cohort [NFBC] 1966 and NFBC1986). Here we identify intronic genetic variants within the *TKT, ARGHAP42,* and *ADAMTS13* genes and intergenic upstream (5’) of the *B3GALT5* and *SSBP2* genes that are associated with prolonged gestation.

## Results

### Discovery Phase

#### NFBC1966 Variants Associated with post-term birth

Analysis of the NFBC1966 post-term cohort identified six GWAS peaks that were suggestive of global significance (p < 1x10^−5^, Figure 1a and 2a). Two clusters of variants were located within introns in the *B3GALT5* (lead SNP rs1534080) and *DNHD1* (rs12285957) genes. An additional four clusters of variants were located within intergenic regions on chromosomes 10 (2 regions), 12, and 15 (Table 1).

**Figure 1.**
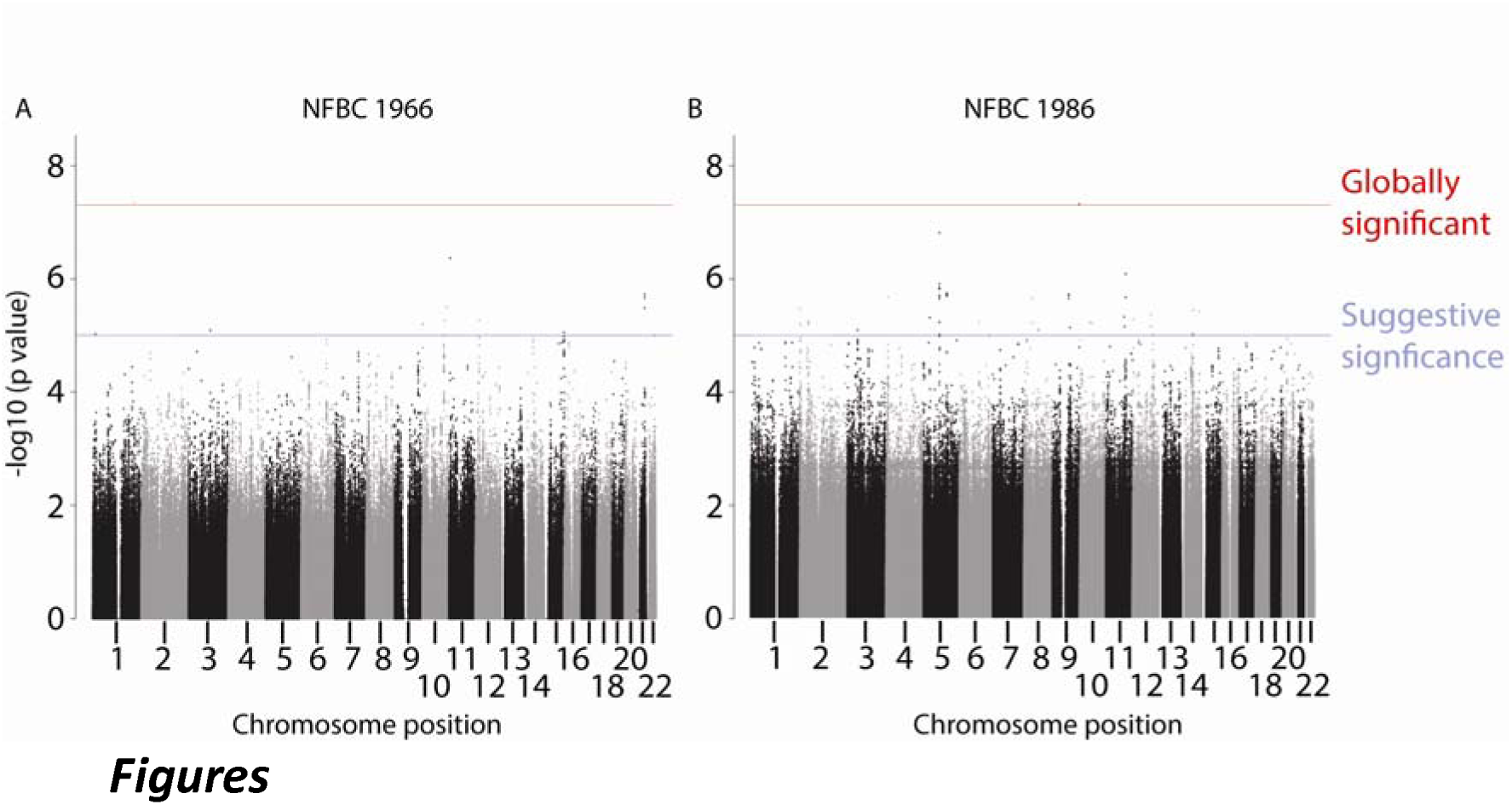
Manhattan plots of the discovery phase of the post-term GWAS for the (A) NFBC1966 and (B) NFBC1986 cohorts. The −log10 observed p-values (2-tailed) for the GWA (y-axis) are plotted versus the chromosomal position of each SNP (x-axis). The blue line indicates significance for followup (p<x10^−5^) through cross-validation, while the red line indicates global significance (5x10-8). Only SNPs in the ADAMTS13 locus in the NFBC1986 cohort reach genome-wide significance in the discovery phase.

**Figure 2.**
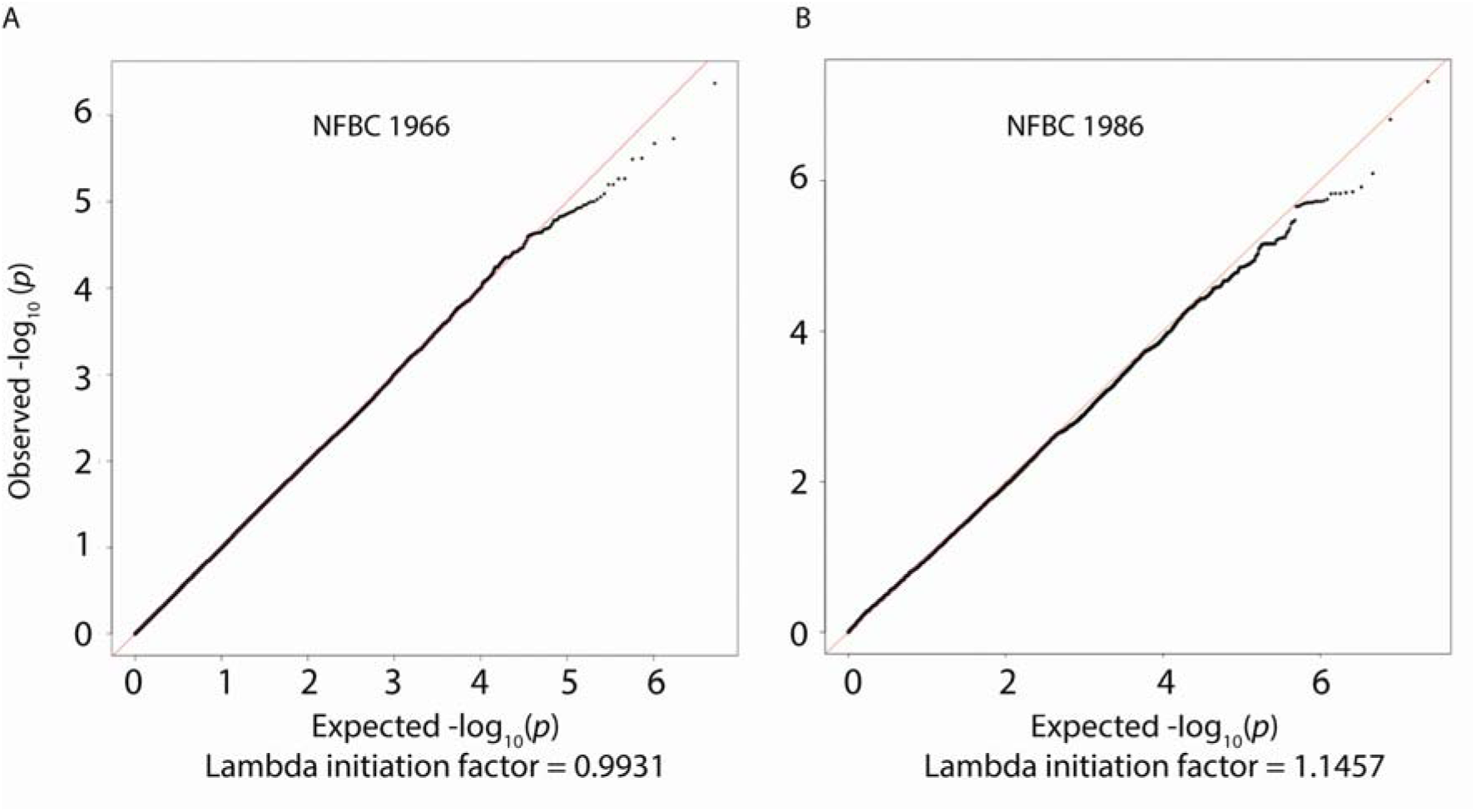
Q-Q plots of the of the quantiles of expected versus observed −log10(p-value) of the association with gestational age in the (A) 1966 and (B) 1986 cohort. The negative logarithm of the expected (x-axis) and the observed (y-axis) p-values for the GWA analysis is plotted for each SNP (black dots). Deviation from the red line indicates points whose observed values are deviating from the null hypothesis of no true association. Inflation factors (λ) near 1 suggest that population stratification was adequately controlled.

**Table 1.**
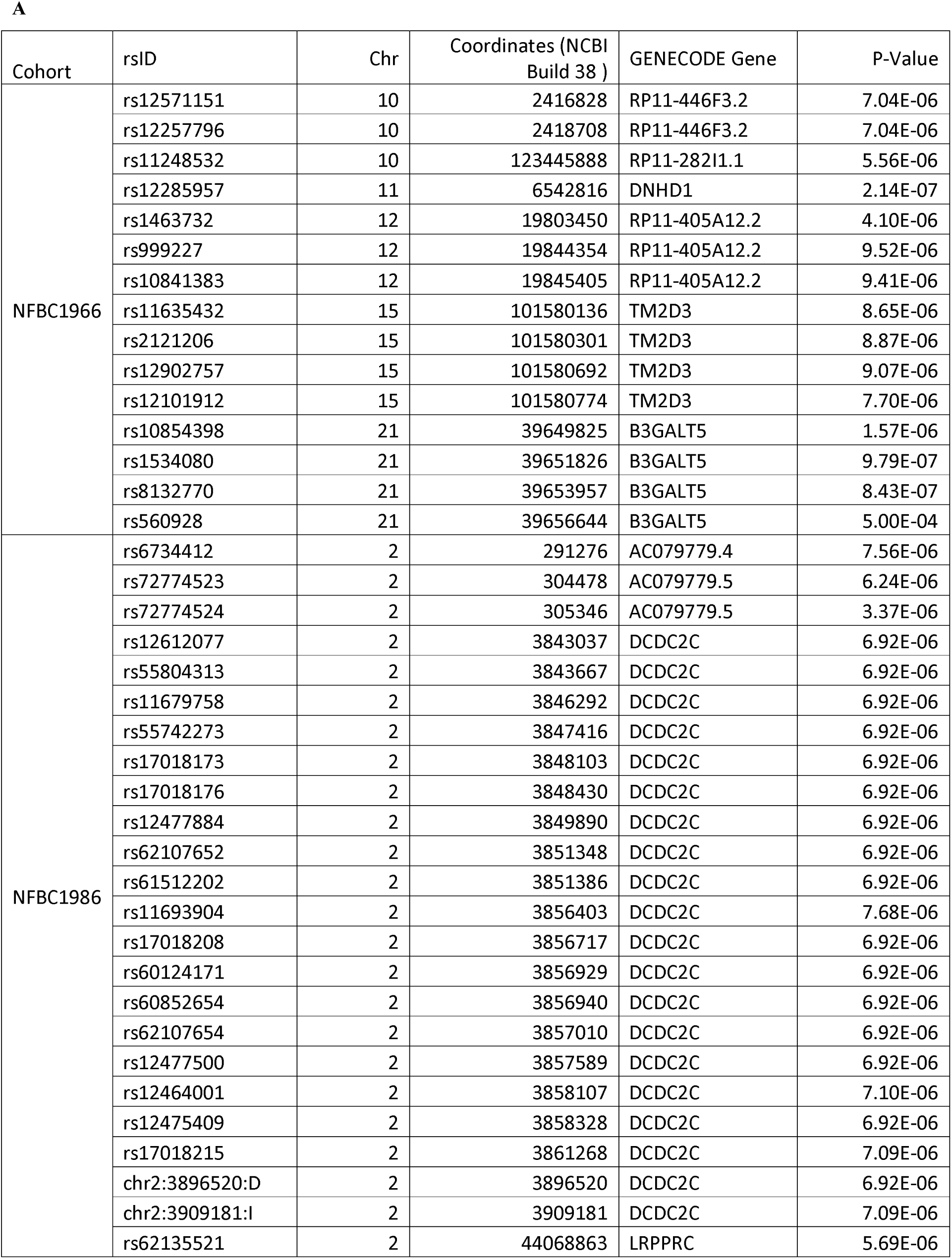
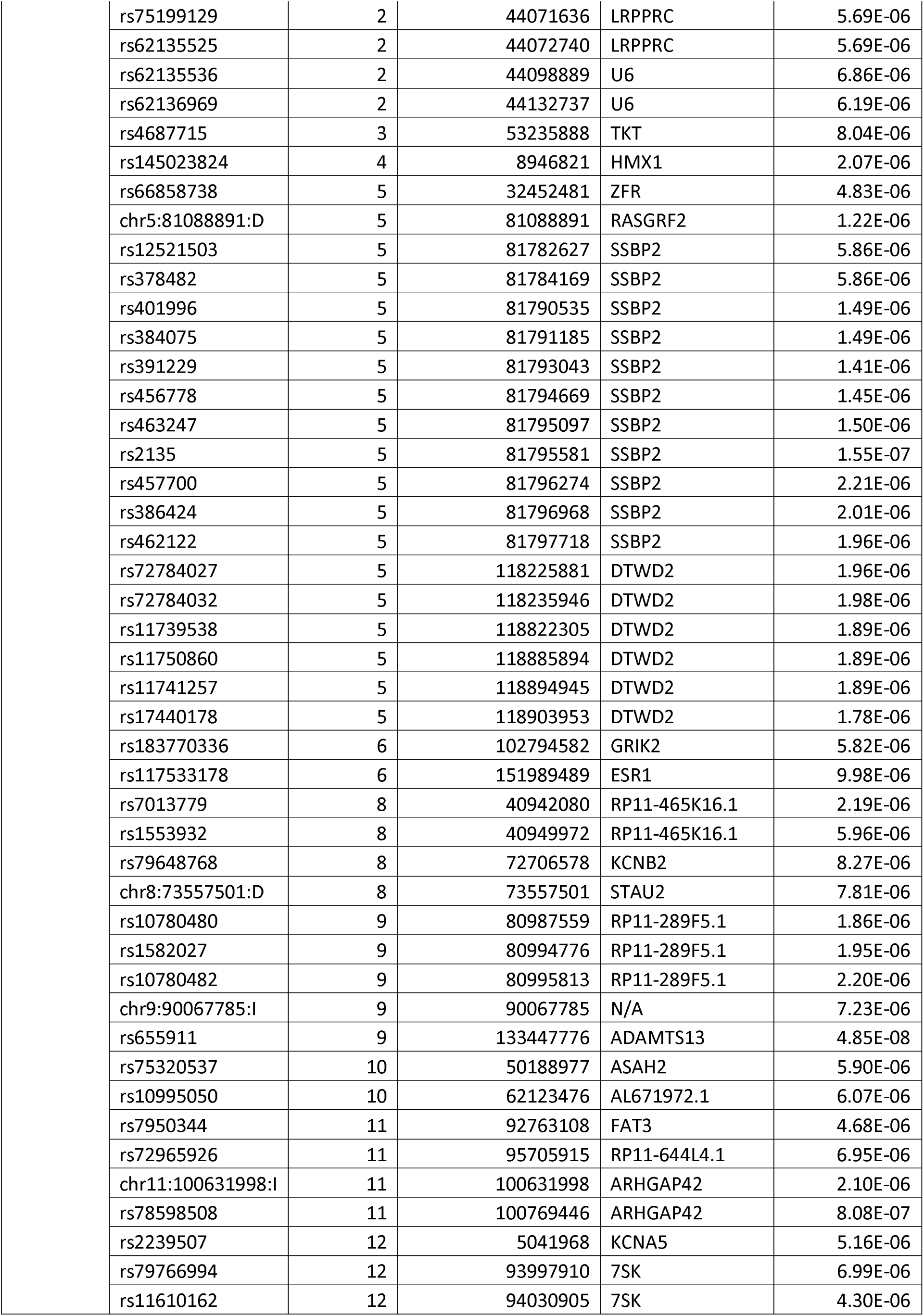
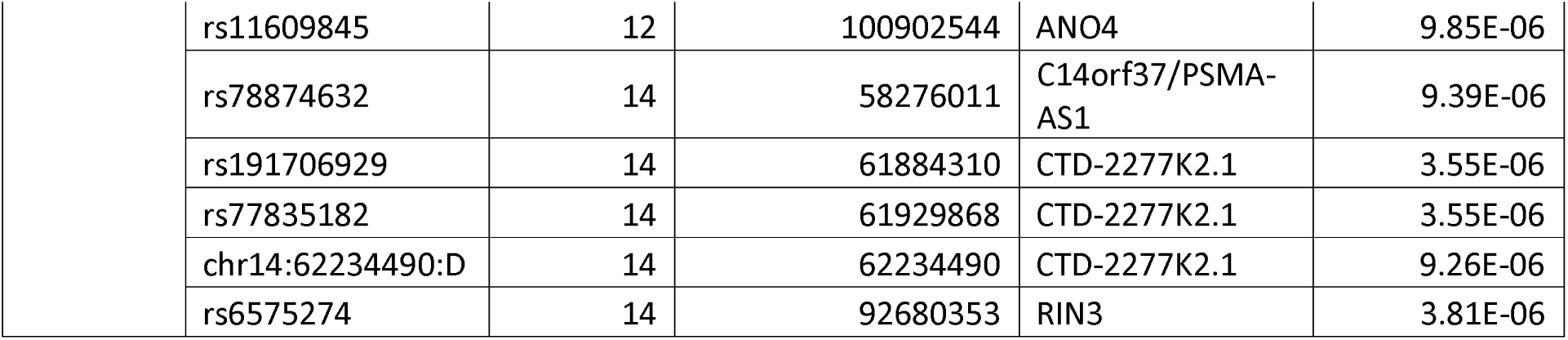
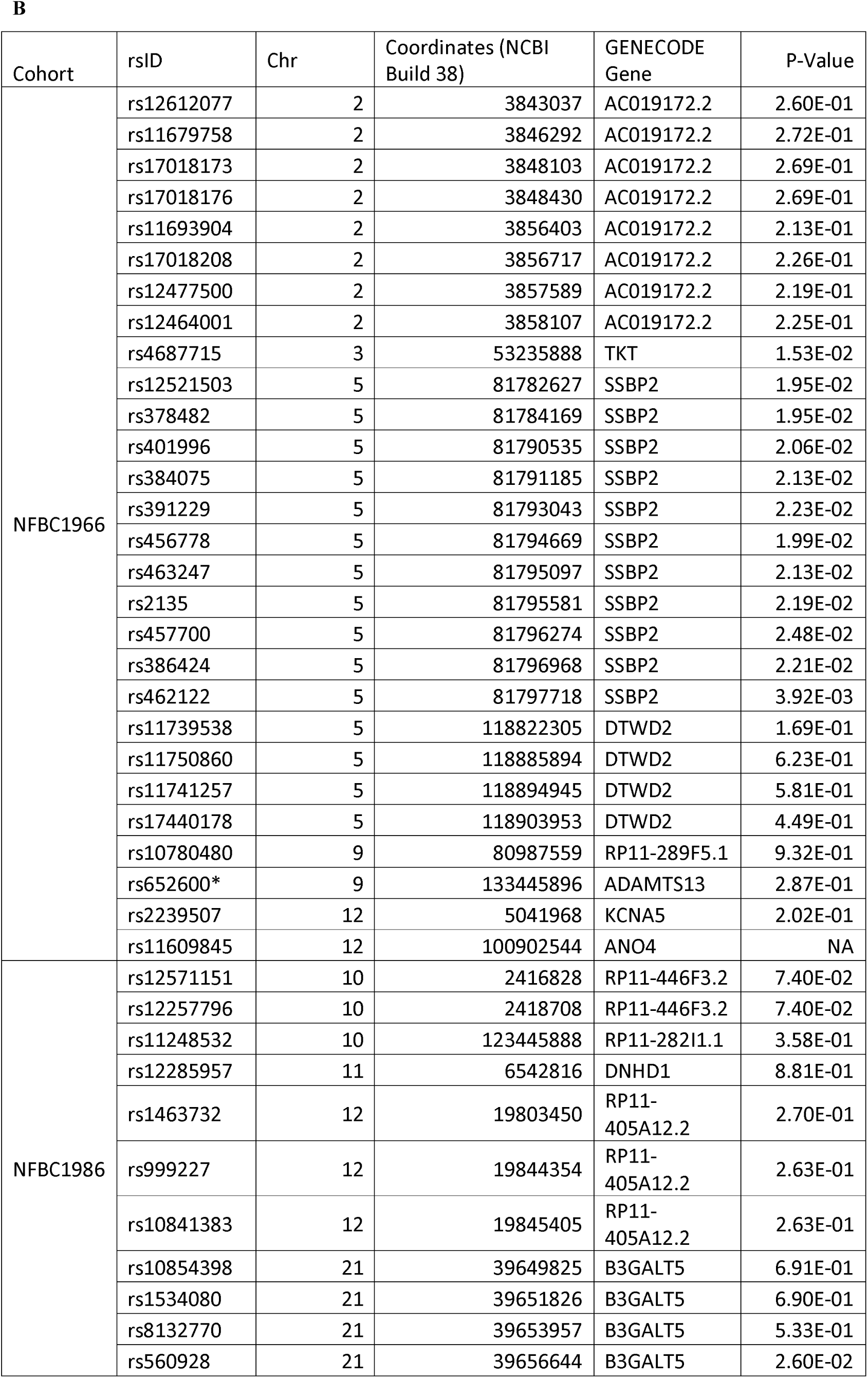
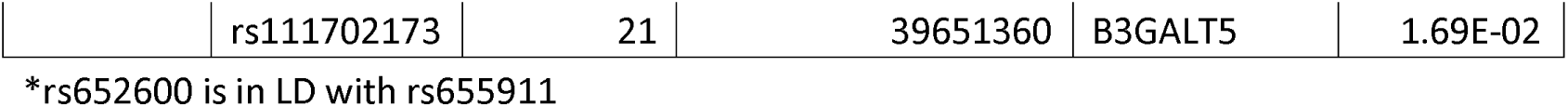
Cross-Validation of the NFBC1966 and NFBC1986 cohorts resulted in five significant loci: B3GALT5, SSBP2, TKT, ARGHAP42, and ADAMTS13. Using a cross-validation methodology of a discovery phase (A, GWAS p < 1x10^−5^) and a validation phase (B, GWAS p<0.05), it was found that the B3GALT5, SSBP2, TKT, and ARGHAP42 loci are significantly associated with post-term birth. Additionally, the *ADAMTS13* locus reached global significance (p < 5x10^−8^).

| Cohort | rsID | Chr | Coordinates (NCBI Build 38 ) | GENECODE Gene | P-Value |
| --- | --- | --- | --- | --- | --- |
| NFBC1966 | rs12571151 | 10 | 2416828 | RP11-446F3.2 | 7.04E-06 |
|  | rs12257796 | 10 | 2418708 | RP11-446F3.2 | 7.04E-06 |
|  | rs11248532 | 10 | 123445888 | RP11-282I1.1 | 5.56E-06 |
|  | rs12285957 | 11 | 6542816 | DNHD1 | 2.14E-07 |
|  | rs1463732 | 12 | 19803450 | RP11-405A12.2 | 4.10E-06 |
|  | rs999227 | 12 | 19844354 | RP11-405A12.2 | 9.52E-06 |
|  | rs10841383 | 12 | 19845405 | RP11-405A12.2 | 9.41E-06 |
|  | rs11635432 | 15 | 101580136 | TM2D3 | 8.65E-06 |
|  | rs2121206 | 15 | 101580301 | TM2D3 | 8.87E-06 |
|  | rs12902757 | 15 | 101580692 | TM2D3 | 9.07E-06 |
|  | rs12101912 | 15 | 101580774 | TM2D3 | 7.70E-06 |
|  | rs10854398 | 21 | 39649825 | B3GALT5 | 1.57E-06 |
|  | rs1534080 | 21 | 39651826 | B3GALT5 | 9.79E-07 |
|  | rs8132770 | 21 | 39653957 | B3GALT5 | 8.43E-07 |
|  | rs560928 | 21 | 39656644 | B3GALT5 | 5.00E-04 |
| NFBC1986 | rs6734412 | 2 | 291276 | AC079779.4 | 7.56E-06 |
|  | rs72774523 | 2 | 304478 | AC079779.5 | 6.24E-06 |
|  | rs72774524 | 2 | 305346 | AC079779.5 | 3.37E-06 |
|  | rs12612077 | 2 | 3843037 | DCDC2C | 6.92E-06 |
|  | rs55804313 | 2 | 3843667 | DCDC2C | 6.92E-06 |
|  | rs11679758 | 2 | 3846292 | DCDC2C | 6.92E-06 |
|  | rs55742273 | 2 | 3847416 | DCDC2C | 6.92E-06 |
|  | rs17018173 | 2 | 3848103 | DCDC2C | 6.92E-06 |
|  | rs17018176 | 2 | 3848430 | DCDC2C | 6.92E-06 |
|  | rs12477884 | 2 | 3849890 | DCDC2C | 6.92E-06 |
|  | rs62107652 | 2 | 3851348 | DCDC2C | 6.92E-06 |
|  | rs61512202 | 2 | 3851386 | DCDC2C | 6.92E-06 |
|  | rs11693904 | 2 | 3856403 | DCDC2C | 7.68E-06 |
|  | rs17018208 | 2 | 3856717 | DCDC2C | 6.92E-06 |
|  | rs60124171 | 2 | 3856929 | DCDC2C | 6.92E-06 |
|  | rs60852654 | 2 | 3856940 | DCDC2C | 6.92E-06 |
|  | rs62107654 | 2 | 3857010 | DCDC2C | 6.92E-06 |
|  | rs12477500 | 2 | 3857589 | DCDC2C | 6.92E-06 |
|  | rs12464001 | 2 | 3858107 | DCDC2C | 7.10E-06 |
|  | rs12475409 | 2 | 3858328 | DCDC2C | 6.92E-06 |
|  | rs17018215 | 2 | 3861268 | DCDC2C | 7.09E-06 |
|  | chr2:3896520:D | 2 | 3896520 | DCDC2C | 6.92E-06 |
|  | chr2:3909181:I | 2 | 3909181 | DCDC2C | 7.09E-06 |
|  | rs62135521 | 2 | 44068863 | LRPPRC | 5.69E-06 |

|  |  |  |  |  |
| --- | --- | --- | --- | --- |
| rs75199129 | 2 | 44071636 | LRPPRC | 5.69E-06 |
| rs62135525 | 2 | 44072740 | LRPPRC | 5.69E-06 |
| rs62135536 | 2 | 44098889 | U6 | 6.86E-06 |
| rs62136969 | 2 | 44132737 | U6 | 6.19E-06 |
| rs4687715 | 3 | 53235888 | TKT | 8.04E-06 |
| rs145023824 | 4 | 8946821 | HMX1 | 2.07E-06 |
| rs66858738 | 5 | 32452481 | ZFR | 4.83E-06 |
| chr5:81088891:D | 5 | 81088891 | RASGRF2 | 1.22E-06 |
| rs12521503 | 5 | 81782627 | SSBP2 | 5.86E-06 |
| rs378482 | 5 | 81784169 | SSBP2 | 5.86E-06 |
| rs401996 | 5 | 81790535 | SSBP2 | 1.49E-06 |
| rs384075 | 5 | 81791185 | SSBP2 | 1.49E-06 |
| rs391229 | 5 | 81793043 | SSBP2 | 1.41E-06 |
| rs456778 | 5 | 81794669 | SSBP2 | 1.45E-06 |
| rs463247 | 5 | 81795097 | SSBP2 | 1.50E-06 |
| rs2135 | 5 | 81795581 | SSBP2 | 1.55E-07 |
| rs457700 | 5 | 81796274 | SSBP2 | 2.21E-06 |
| rs386424 | 5 | 81796968 | SSBP2 | 2.01E-06 |
| rs462122 | 5 | 81797718 | SSBP2 | 1.96E-06 |
| rs72784027 | 5 | 118225881 | DTWD2 | 1.96E-06 |
| rs72784032 | 5 | 118235946 | DTWD2 | 1.98E-06 |
| rs11739538 | 5 | 118822305 | DTWD2 | 1.89E-06 |
| rs11750860 | 5 | 118885894 | DTWD2 | 1.89E-06 |
| rs11741257 | 5 | 118894945 | DTWD2 | 1.89E-06 |
| rs17440178 | 5 | 118903953 | DTWD2 | 1.78E-06 |
| rs183770336 | 6 | 102794582 | GRIK2 | 5.82E-06 |
| rs117533178 | 6 | 151989489 | ESR1 | 9.98E-06 |
| rs7013779 | 8 | 40942080 | RP11-465K16.1 | 2.19E-06 |
| rs1553932 | 8 | 40949972 | RP11-465K16.1 | 5.96E-06 |
| rs79648768 | 8 | 72706578 | KCNB2 | 8.27E-06 |
| chr8:73557501:D | 8 | 73557501 | STAU2 | 7.81E-06 |
| rs10780480 | 9 | 80987559 | RP11-289F5.1 | 1.86E-06 |
| rs1582027 | 9 | 80994776 | RP11-289F5.1 | 1.95E-06 |
| rs10780482 | 9 | 80995813 | RP11-289F5.1 | 2.20E-06 |
| chr9:90067785:I | 9 | 90067785 | N/A | 7.23E-06 |
| rs655911 | 9 | 133447776 | ADAMTS13 | 4.85E-08 |
| rs75320537 | 10 | 50188977 | ASAH2 | 5.90E-06 |
| rs10995050 | 10 | 62123476 | AL671972.1 | 6.07E-06 |
| rs7950344 | 11 | 92763108 | FAT3 | 4.68E-06 |
| rs72965926 | 11 | 95705915 | RP11-644L4.1 | 6.95E-06 |
| chr11:100631998:I | 11 | 100631998 | ARHGAP42 | 2.10E-06 |
| rs78598508 | 11 | 100769446 | ARHGAP42 | 8.08E-07 |
| rs2239507 | 12 | 5041968 | KCNA5 | 5.16E-06 |
| rs79766994 | 12 | 93997910 | 7SK | 6.99E-06 |
| rs11610162 | 12 | 94030905 | 7SK | 4.30E-06 |
| rs11609845 | 12 | 100902544 | ANO4 | 9.85E-06 |
| rs78874632 | 14 | 58276011 | C14orf37/PSMA-AS1 | 9.39E-06 |
| rs191706929 | 14 | 61884310 | CTD-2277K2.1 | 3.55E-06 |
| rs77835182 | 14 | 61929868 | CTD-2277K2.1 | 3.55E-06 |
| chr14:62234490:D | 14 | 62234490 | CTD-2277K2.1 | 9.26E-06 |
| rs6575274 | 14 | 92680353 | RIN3 | 3.81E-06 |

**Table 1. Cross-Validation of the NFBC1966 and NFBC1986 cohorts resulted in five significant loci: B3GALT5, SSBP2, TKT, ARGHAP42, and ADAMTS13.**
| Cohort | rsID | Chr | Coordinates (NCBI Build 38) | GENECODE Gene | P-Value |
| --- | --- | --- | --- | --- | --- |
| NFBC1966 | rs12612077 | 2 | 3843037 | AC019172.2 | 2.60E-01 |
|  | rs11679758 | 2 | 3846292 | AC019172.2 | 2.72E-01 |
|  | rs17018173 | 2 | 3848103 | AC019172.2 | 2.69E-01 |
|  | rs17018176 | 2 | 3848430 | AC019172.2 | 2.69E-01 |
|  | rs11693904 | 2 | 3856403 | AC019172.2 | 2.13E-01 |
|  | rs17018208 | 2 | 3856717 | AC019172.2 | 2.26E-01 |
|  | rs12477500 | 2 | 3857589 | AC019172.2 | 2.19E-01 |
|  | rs12464001 | 2 | 3858107 | AC019172.2 | 2.25E-01 |
|  | rs4687715 | 3 | 53235888 | TKT | 1.53E-02 |
|  | rs12521503 | 5 | 81782627 | SSBP2 | 1.95E-02 |
|  | rs378482 | 5 | 81784169 | SSBP2 | 1.95E-02 |
|  | rs401996 | 5 | 81790535 | SSBP2 | 2.06E-02 |
|  | rs384075 | 5 | 81791185 | SSBP2 | 2.13E-02 |
|  | rs391229 | 5 | 81793043 | SSBP2 | 2.23E-02 |
|  | rs456778 | 5 | 81794669 | SSBP2 | 1.99E-02 |
|  | rs463247 | 5 | 81795097 | SSBP2 | 2.13E-02 |
|  | rs2135 | 5 | 81795581 | SSBP2 | 2.19E-02 |
|  | rs457700 | 5 | 81796274 | SSBP2 | 2.48E-02 |
|  | rs386424 | 5 | 81796968 | SSBP2 | 2.21E-02 |
|  | rs462122 | 5 | 81797718 | SSBP2 | 3.92E-03 |
|  | rs11739538 | 5 | 118822305 | DTWD2 | 1.69E-01 |
|  | rs11750860 | 5 | 118885894 | DTWD2 | 6.23E-01 |
|  | rs11741257 | 5 | 118894945 | DTWD2 | 5.81E-01 |
|  | rs17440178 | 5 | 118903953 | DTWD2 | 4.49E-01 |
|  | rs10780480 | 9 | 80987559 | RP11-289F5.1 | 9.32E-01 |
|  | rs652600* | 9 | 133445896 | ADAMTS13 | 2.87E-01 |
|  | rs2239507 | 12 | 5041968 | KCNA5 | 2.02E-01 |
|  | rs11609845 | 12 | 100902544 | ANO4 | NA |
| NFBC1986 | rs12571151 | 10 | 2416828 | RP11-446F3.2 | 7.40E-02 |
|  | rs12257796 | 10 | 2418708 | RP11-446F3.2 | 7.40E-02 |
|  | rs11248532 | 10 | 123445888 | RP11-282I1.1 | 3.58E-01 |
|  | rs12285957 | 11 | 6542816 | DNHD1 | 8.81E-01 |
|  | rs1463732 | 12 | 19803450 | RP11-405A12.2 | 2.70E-01 |
|  | rs999227 | 12 | 19844354 | RP11-405A12.2 | 2.63E-01 |
|  | rs10841383 | 12 | 19845405 | RP11-405A12.2 | 2.63E-01 |
|  | rs10854398 | 21 | 39649825 | B3GALT5 | 6.91E-01 |
|  | rs1534080 | 21 | 39651826 | B3GALT5 | 6.90E-01 |
|  | rs8132770 | 21 | 39653957 | B3GALT5 | 5.33E-01 |
|  | rs560928 | 21 | 39656644 | B3GALT5 | 2.60E-02 |
|  | rs111702173 | 21 | 39651360 | B3GALT5 | 1.69E-02 |

\*rs652600 is in LD with rs655911

#### NFBC1986 Variants Associated with post-term birth

Analysis of the NFBC1986 post-term cohort identified twenty-five significant GWAS peaks (p < 1x10^−5^, Table 1, Figure 1b and 2b). The lead SNPs for fourteen of these GWAS peaks were intronic: AC079779.5 (rs72774524), DCDC2C (rs12612077), TKT (rs4687715), DTWD2 (rs17440178), ESR1 (rs117533178), KCNB2 (rs79648768), ADAMTS13 (rs655911), ASAH2 (rs75320537), FAT3 (rs7950344), ARHGAP42 (rs78598508), AN04 (rs11609845), C14orf37/PSMA-AS1 (rs78874632), CTD-2277K2.1 (rs191706929), and RIN3 (rs6575274). Only the SNP at the ADAMTS13 locus was globally significant (p < 5x10^−8^).

Eleven intergenic loci were also associated with gestational age in the NFBC1986 cohort. These intergenic loci were located ≤116kbp from a coding exon (gene): LRPPRC (rs62135521, 73 kb upstream), HMX1 (rs145023824, 75 kb upstream), ZFR (rs66858738, 7.7 kb upstream), SSBP2 (rs2135, 31 kb upstream), GRIK2 (rs183770336, 724 kb downstream), RP11-465K16.1 (rs7013779, 40 kb upstream), RP11-289F5.1 (rs10780480, 116 kb upstream), AL671972.1 (rs10995050, 7.2 kb downstream), RP11-644L4.1 (rs72965926, 5.3 kb downstream), KCNA5 (rs2239507, 2 kb upstream), and 7SK (rs11610162, 11 kb upstream)

Of the list of genes located close to the loci that were significantly associated with post-term birth in the 1986 cohort, only ARHGAP42 has previously been associated with a developmental phenotype (age at menarche in a Japanese population, rs12800752)[21].

### Validation of Results of the Discovery Phase

The six NFBC1966 loci and twenty-five NFBC1986 loci were tested for cross-validation (*i.e.* significance) in the opposite cohort. Of these loci, none were associated with gestational age at a p value of ≤1x10^−5^ in both cohorts. However, the *B3GALT5, SSBP2,* and *TKT* GWAS loci (hereafter referred to as post-term loci) were validated as significant in both the 1966 and 1986 cohorts (*i.e.* discovery p < 1x10^−5^ and validation p<0.05, Tables 1A and 1B). *ADAMTS13* reached global significance (p < 5x10^−8^ [22]) in the NFBC1986 cohort, and was included in the post-term loci for further analyses (Table 1B).

The rs78598508 variant, which is located within *ARHGAP42* and associated with gestational age in the 1986 cohort, was not measured in the 1966 cohort. Furthermore, rs78598508 was not in strong LD (> 0.9 r^2^) with any other variants. Therefore, rs78598508 could not be tested for cross-validation in this study (Table 1).

### Identification of Spatial and Functional Connections to Post-Term Birth

#### Spatial Associations

The post-term associated SNPs we identified fall outside of known coding regions and, as such, there is no *a priori* reason to assume impact through direct disruption of protein function. Therefore, the effect of these SNPs can be better explained through a model of gene regulation that incorporates the hypothesis that long distance SNP-Gene interactions make a significant contribution to genetically determined phenotypes[23, 24]. Observations of long distance SNP-gene regulatory interactions in diabetes (variants in the FTO gene alter the expression of the IRX3 gene)[25], inflammatory bowel disease (variants strongly interact with an anti-inflammatory gene located 380 kb away, and not closer anti-inflammatory genes) [26], and growth[27] are consistent with this hypothesis.

We screened Haploreg v4.1 (1000 genomes haplotype data), to show that none of the lead SNPs in the *B3GALT5, ADAMTS13, SSBP2, ARHGAP42,* and *TKT* loci are in linkage disequilibrium (LD, r^2^ > 0.95 and D’ > 0.95) with any variants located within exons or critical transcriptional processing sequences (*e.g.* intronic branch sites, polyA signals, or transcription termination signals). Thus, there is no evidence that these SNPs directly impact on protein function through aberrant transcript processing. As such, we hypothesized that the post-term SNPs were affecting enhancer regions (*i.e.* short genomic regions that are bound by transcription factors) and altering the transcriptional regulation of distant genes.

Interactions between the lead SNPs at each post-term locus and distant genes were screened for using GWAS3D (Figure 3, Table 2). The *B3GALT5* locus spatially connects to the Down Syndrome Cell Adhesion Molecule *(DSCAM)* locus. *DSCAM* is a member of the immunoglobulin superfamily of cell adhesion molecules (Ig-CAMs) that are involved in human central and peripheral nervous system development. The *ADAMTS13* locus spatially connects to the *SLC2A6* (9q34), *COL5A1* (9q34.2-q34.3), and *RABGAP1L* (1q24) loci. This is notable as *ADAMTS13, SLC2A6, COL5A1,* and additional genes within 9q34 have been implicated in the coagulation process and associated with ovarian function[28]. The *TKT* locus spatially connects to an intergenic region in 21p11.2 which contains predicted open reading frames that encode undefined proteins.

**Figure 3.**
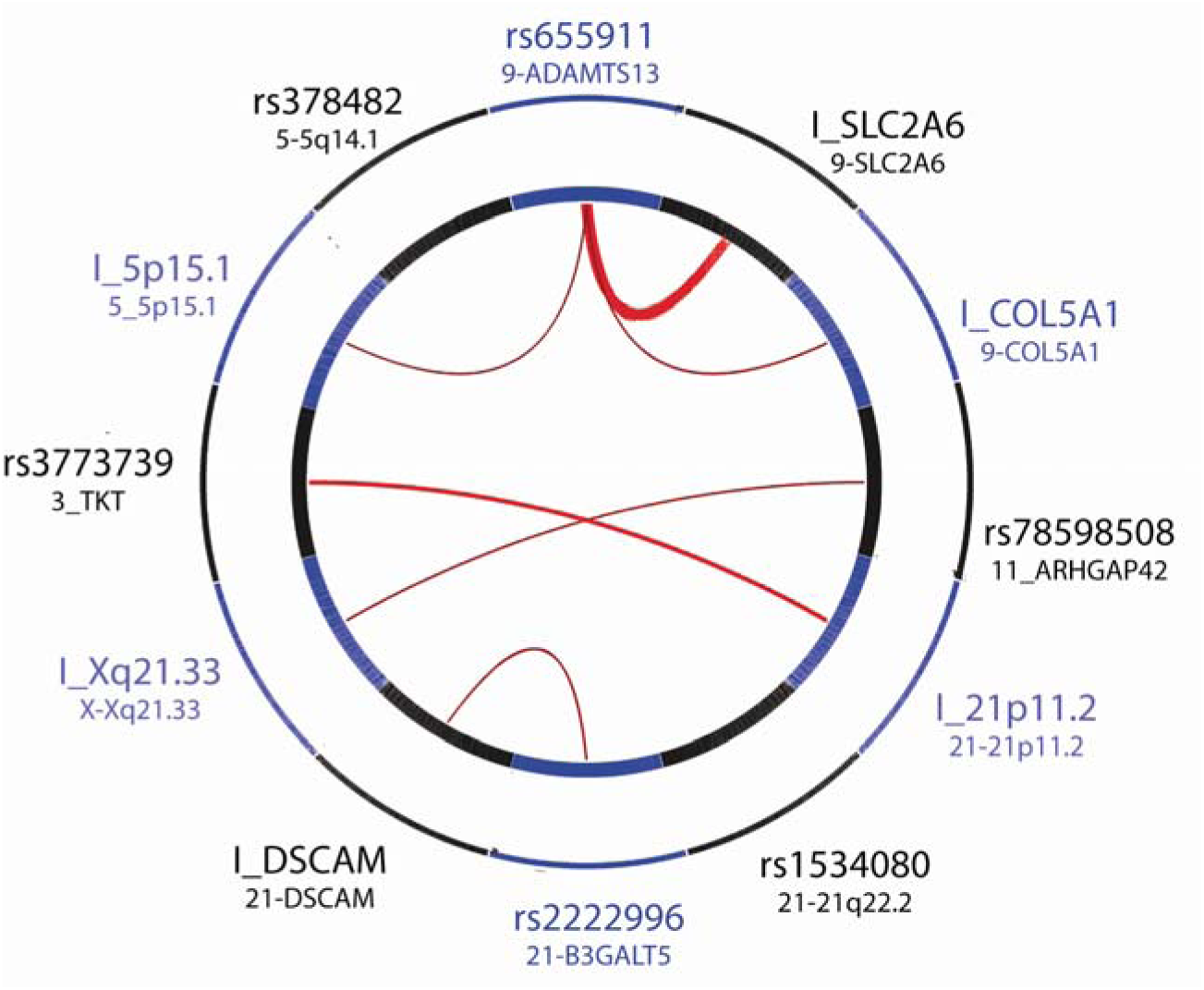
Spatial Results from GWAS3D identify 4 significant spatial connections between loci in the validated GWAS data and distant genomic regions. The SNP associated with the *ADAMTS13* locus had multiple spatial connections, *SSBP2* had none, while the others only exhibited a single spatial association. Only the spatial connections with high confidence scores are plotted here (thickness of the red line).

**Table 2.** Spatial Results from GWAS3D identify significant spatial connections between loci in the validated GWAS data and distant genomic regions. The *ADAMTS13* locus had multiple spatial connections, *SSBP2* had none, while the others only exhibited a single spatial association.

| SNP | Locus | Spatial Connections |
| --- | --- | --- |
| rs560928 | B3GALT5 | chr21:42080001-42090000 |
| rs2135 | SSBP2 | None |
| rs4687715 | TKT | chr21:9650001-9660000 |
| rs655911 | ADAMTS13 | chr9:136340001-136350000, chr9:136330001-136340000, chr9:137560001-137570000, chr5:18160001-18170000, chr1:174900001-174910000 |
| rs78598508 | ARHGAP42 | chrX:93780001-93790000 |
| rs111702173 | B3GALT5 | chr21:42080001-42090000 |

#### Locus-Specific Associations with Gene Expression

Functional-regulatory roles for the SNPs we identified in this study were refined by testing the spatial SNP-gene pairs for significant eQTLs using the GTEx database (version 6). Variants in the *ADAMTS13* (Tibial Nerve Tissue, p=1.50x10^−8^), *B3GALT5* (Thyroid Tissue, p=9.0x10^−5^), and *TKT* (Left Ventricular Heart Tissue, p=2.0x10^−5^) loci associate with altered expression changes within these genes, confirming that these SNPs fall within loci that regulate their local gene landscape (Table 3). In addition, rs655911 (intronic, *ADAMTS13*) also showed eQTL associations with the expression of *SLC2A6,* reinforcing the significance of the spatial connection. The variant associated with the *ARHGAP42* locus (rs78598508) showed no significant eQTLs.

**Table 3.**
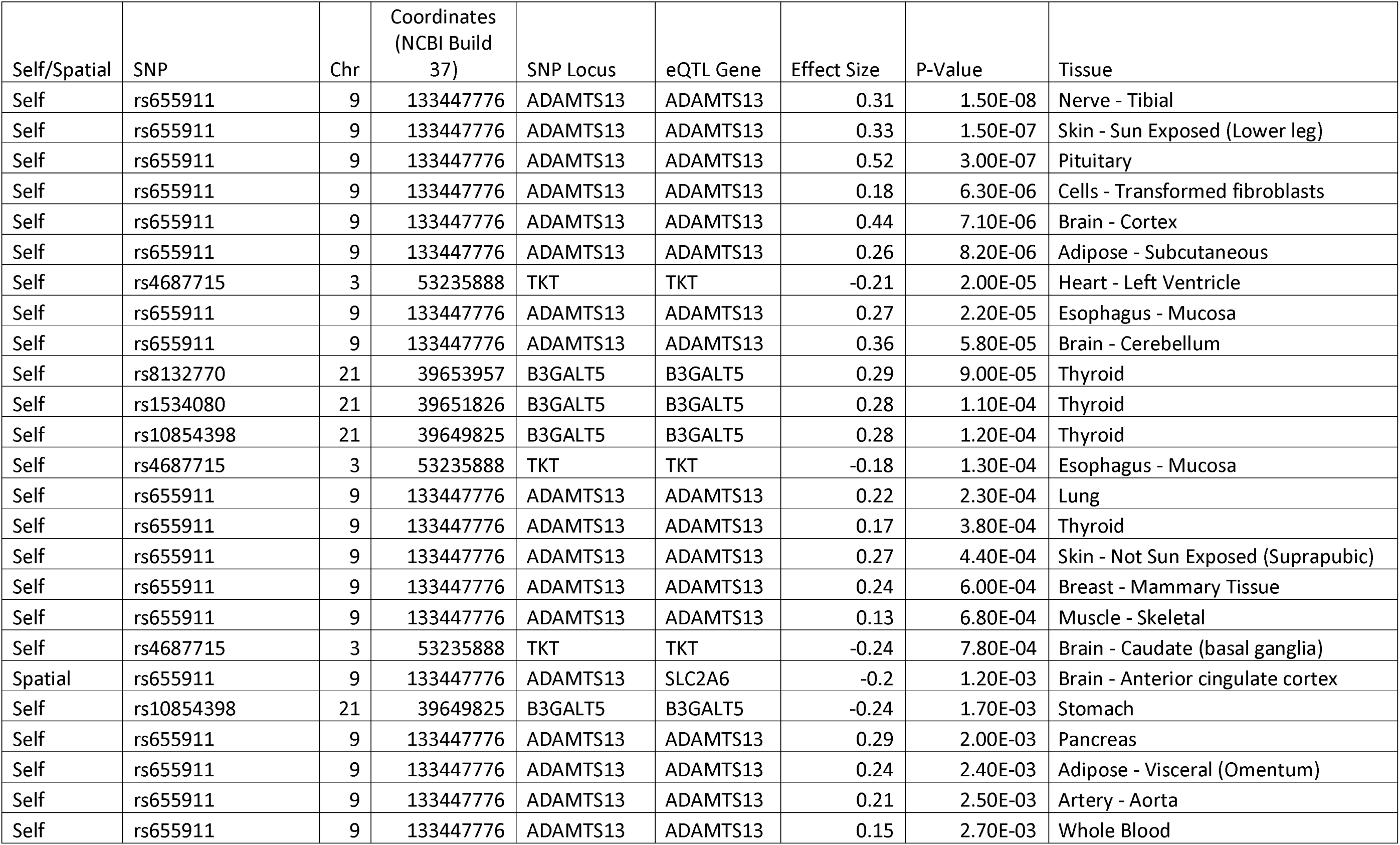
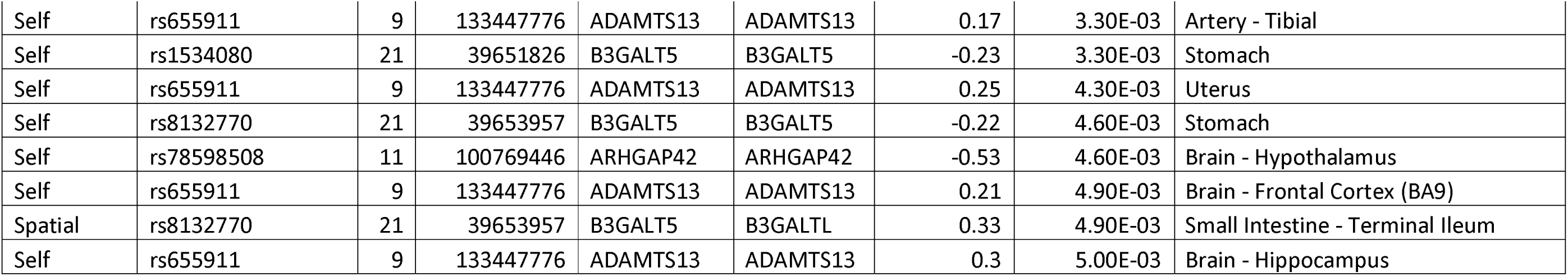
eQTL analysis supports the ADAMTS13-SLC2A6 spatial connection, but also confirms self-eQTLs for the ADAMTS13, TKT, and B3GALT5 post-term loci. Using GTEx to determine effect size and significance of SNP-gene expression associations, it was determined that a number of eQTLs exist that support the spatial connections. This table includes all self- and spatial-eQTLs with p < 5 x 10^−3^ in GTEx (version 6).

The SNP rs2135, which was 44kb upstream of *SSBP2,* did not show any evidence of spatial connections to other regions. Moreover, there was no evidence of a cis-eQTL between this SNPs and the *SSBP2* gene itself. A global survey of eQTL associations with the *SSBP2* SNPs, within GTEx, did not identify any globally significant eQTLs (Supplemental Figure 1). Analyses indicated putative eQTLs between the SSBP2 lead SNP (rs2135; 5q14.1) and: HBG1 (11p15.5, Lung, 1.10x10^−5^); HLA-DRB5 (6p21.3, Whole Blood, 2.50x10^−5^); and FYB (5p13.1, Mucosa of the Esophagus, 4.00x10^−5^) (Supplemental Figure 1).

#### Locus-Specific Associations with Gene Expression

The lead SNPs that were proximal to *B3GALT5* (rs11702173, rs560928), *TKT* (rs4687715), *SSBP2* (rs2135), and *ARHGAP42* (rs78598508) were screened for enhancer activity (Figure 4). *ADAMTS13* (rs655911) was unable to be cloned and could not be tested. Cloning loci with alternate alleles allowed the measurement of the effect of genetic variation on the observed enhancer activity. Luciferase assays in Hela cells revealed a pronounced enhancer effect for the loci containing rs4687715, rs111702173, rs78598508, and rs560928 and a repressive effect for rs2135 (Figure 4). For rs4687715 and rs560928, the region shows a differential enhancer allelic effect. Therefore, for two of the five loci tested, the enhancer activity associated with these gestational age associated regions is sensitive to the identity of the haplotype at the SNP position. For the remaining three regions, the SNP tested did not have a measurable allelic effect in HeLa cells, but still showed significant enhancer/insulator capabilities.

**Figure 4.**
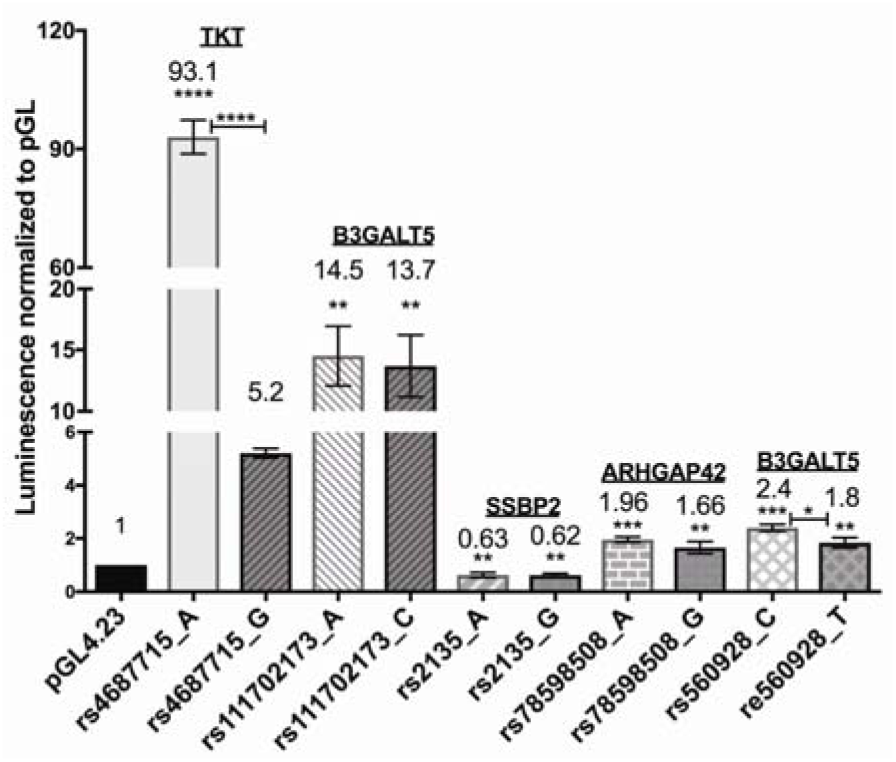
Post-term associated SNPs show allele specific enhancer and repressor effects. All amplified regions, except that containing rs21355, acted as enhancers. There were significant differences (p<0.00001) between the enhancer activity of the ‘A’ and ‘G’ versions of rs4687715. Similarly, there were significant (p<0.05) differences between the enhancer activity of the ‘C’ and ‘T’ alleles of rs560928 in HeLa cells. Notably, DNA amplicons containing the A and G alleles of rs2135 acted as a repressor of basal activity. PCR-amplified genomic DNA with the indicated SNP variants were assayed for their ability to drive luciferase expression in HeLa cells. The increase in luminescence indicates that competence for transcription depends on the alellic version of these SNPs. Error bars represent ±SEM from four biological replicates and significance was determined by One-Way ANOVA. Asterisks above the bars denote significant differences compared to the empty pGL4.23 vector control. Allele specific differences that are significant are indicated. (^****^p<0.0001, ^***^p<0.001, ^**^p0.01, ^*^p<0.05).

## Discussion

Previously, there has been indirect evidence of a genetic component to post-term birth, as children born post-term are more likely to have a sibling or mother born post-term[6]. This study is the first to identify specific genetic variants in proximity to the *B3GALT5, ADAMTS13, SSBP2,* and *TKT* genes as being associated with prolonged gestation. Data on the spatial connections with these loci, eQTLs and enhancer activity is consistent with these post-term variants acting as functional determinants of gestational length. Thus, the SNPs associated with the *B3GALT5, ADAMTS13, SSBP2,* and *TKT* loci may alter the expression of these genes, contributing to the post-term phenotype by affecting regulation of processes involved in human development such as growth and metabolism, and, more specifically, hematopoiesis.

### Post-term Associated TKT and SSBP2 are linked to Alterations in Cellular Growth, Proliferation, and Metabolism

Alterations in cellular growth, proliferation, and metabolism pre-program biological development (*e.g.* pentose phosphate pathway) resulting in an amplified risk of chronic noncommunicable disease[29]. Proteins encoded by the *TKT* and *SSBP2* genes are involved in the cellular growth, proliferation, and metabolism and are thus capable of altering developmental trajectories. For example, TKT is involved in carbohydrate metabolism[30] and could contribute to the later-in-life increased adiposity and risk of metabolic syndrome in children and adults born post-term[6, 7].

### Post-term Associated TKT is linked to Alterations in Cell Cycle and Growth

*TKT* dysregulation has an important role in cellular growth rates. The *TKT* gene encodes a protein that contributes to the main carbohydrate metabolic pathways by connecting the pentose phosphate pathway (PPP) to glycolysis. This process results in NADPH synthesis. NADPH is part of the control for reactive oxygen species, which were found to be imbalanced in post-term births[31]. A critical role for TKT (highly expressed in oocytes) is in oocyte cell cycle progression and maturation[30]. Under-expression of *TKT* in maternal mice contributes to pregnancy resulting in fewer progeny, retarded postnatal growth, and reduced levels of adipose tissue in offspring[32]. This phenotype has obvious similarities to post-term infants, who are typically born lean[14]. Collectively, the effects of aberrant *TKT* expression can be interpreted as being consistent with TKT variation in humans contributing to aberrant gestational timing. Moreover, the links between variation in TKT function and metabolism may help to partially explain the observed links between post-term birth and the later development of symptoms of the metabolic syndrome [6].

### Post-term associated SSBP2 and ADAMTS13 are linked with hematopoiesis and blood disorders

Alterations in *SSBP2* and *ADAMTS13* levels could be affecting gestation through alterations in hematopoietic pathways. There is a large developmental aspect to hematopoiesis during different phases of gestation. After the second month of human gestation, hematopoiesis occurs primarily in the liver, and by the fourth to fifth month of gestation hematopoiesis switches again, this time to the bone marrow (where it occurs for the duration of life). Maturity of the hematopoietic system occurs late in gestation and tracks with gestational age, with the proportions of fetal hemoglobin decreasing during the progression from preterm - term - post-term (83.93% to 68.59% to 60.03%, respectively) [33]. In post-term births, cord blood collected at birth shows differences in levels of polycythemia (increased concentration of hemoglobin in the blood), erythropoietin levels (increased erythropoiesis), mean corpuscular hemoglobin, red blood cell count, neutrophil count, and monocyte count [34]. And finally, primiparity (a risk factor for post-term birth), post-term delivery, or delivery by emergency caesarean section are all correlated with an increased risk of arterial ischaemic stroke (17–20 per 100 000 live births) in the first year of life [35].

Our eQTL analysis showed that the variants in SSBP2 did not show significant evidence of transcriptional regulatory effects on SSBP2 itself. However, there were putative eQTLs with *HBG1, HLA-DRB5* and *FYB* (Supplemental Figure 1). *HBG1* is a key component of the fetal hemoglobin locus that is normally expressed in the fetal liver, spleen and bone marrow, as a part of the constitution of fetal hemoglobin (HbF)[36]. In adults, the beta-globin locus is only accessible (chromatin open, DNase I hypersensitive) in adult erythroid cells, however HBG1 shows chromatin accessibility in both fetal and adult erythroid cells[36]. HLA-DRB5, is a component of the major histocompatibility complex and thus plays a central role in the immune system, specifically antigen presentation and recognition[37]. The FYB gene encodes a hematopoietic-specific protein involved in platelet activation[38]. In mice, FYB knockout affects platelet function and causes mild thrombocytopenia[39]. Therefore, our results are consistent with rs2135 being involved in dysregulation of genes that contribute to the production and development of fetal blood.

The *ADAMTS13* gene encodes a protease that has previously been shown to disrupt the regulation of platelet thrombosis by cleaving Von Willebrand factor [40]. Critically, ADAMTS13 variants are also known to cause neonatal platelet disorders. These disorders include Upshaw-Schulman syndrome (hemolytic anemia) and blood hyper-coagulation (thrombophilia), the second of which is associated with fetal loss[28]. Thus, the identification here of *ADAMTS13* (rs655911) as significantly associated with post-term birth is consistent with earlier findings of hematopoetic phenotypes associating with post-term birth, and could be identifying a key gene in this cause-effect association with gestational length. Notably, the identification of a significant eQTL between a postterm associated *B3GALT5* SNP and *B3GALTL* expression reinforces the significance of the ADAMTS13 variants (Table 3) [21]. Specifically, *B3GALTL* encodes a beta-1,3 glucosyltransferase that is required to glycosylate the *ADAMTS13* gene product as part of its pre-processing for Von Willebrand factor cleavage [41].

Therefore, the role of alterations in expression of both *SSBP2* and *ADAMTS13* could result in a post-term birth phenotype arising through alterations in hematopoiesis. It should be noted that the *SSBP2* variant was shown to repress transcription in the luciferase assay while the *ADAMTS13* locus wasn’t able to be assessed. In future work, identifying the regulatory role of these regions could help elucidate the cause-and-effect relationships between hematopoiesis and gestational length.

### Post-term Birth Versus the Rise of More Intensive Obstetric Management of Birth

Analysing two cohorts from the same geographical region (Northern Finland) enabled us to control for regional and culture-specific differences in routine management of pregnancies. However, two major confounders remain between the 1966 and 1986 cohorts: 1) technological changes led to better estimation and certainty of gestational age in the 1986 cohort; and 2) management practices led to changes in the incidence of induced labor and post-term birth. Firstly, the uncertainty surrounding gestational age prediction in 1966 raises issues around phenotype definition in this cohort. The impact of this ambiguity on ours study was minimized through the use of a narrow term gestational age range of 38 0/7 to 40 0/7.

There have been significant changes to obstetric management over the last 50 years, with a shift from conservative monitoring of prolonged pregnancies through to the current recommendations to induce women who are beyond 41 weeks gestation[5, 8, 42]. The induction of labor became a therapeutic option between 1966 and 1986. Therefore, the NFBC1986 cohort contains births that would have been post-term if not for induction and/or Caesarean-section. Consistent with this, we observed a reduction in numbers of post-term births from approximately 20% in the 1966 NFBC cohort to less than 5% in the 1986 NFBC cohort. However, while the induction of labour is an improvement in obstetric management, it is possible that these individuals have “post-term potential” and carry genetic risks that were not mitigated by the act of induction. Therefore, we excluded all term-born induced births from the 1986 cohort from the analyses. It should be noted that the fact that we identified genetic variants associated with post-term birth is consistent with the idea of post-term potential and induced children having similar genetic health risks as those born post-term.

We have identified genetic variants in proximity to the *B3GALT5, ADAMTS13, SSBP2,* and *TKT* loci as being associated with post-term birth in two birth cohorts (NFBC1966 and NFBC1986). This finding is consistent with previous observations that suggested there was a genetic component to post-term birth[12, 13, 15–20, 43]. Spatial and mRNA expression analyses further showed provided novel clues about how these loci contribute to the regulation and consequences of post-term birth. This study forms a foundation for a better understanding of the genetic and long term metabolic health risks faced by induced and post-term individuals. Since nearly 20% of births in the NFBC cohort were postterm, the long-term risks for induced individuals who have a previously overlooked post-term potential may be a major issue for current health providers.

## Materials and Methods

### Subjects

We undertook a discovery-replication study of two successive birth cohorts from Northern Finland (*i.e.* NFBC 1966[10] and NFBC1986[11]). Both cohorts were recruited from the two northernmost provinces of Finland (*i.e.* Oulu and Lapland). Each cohort followed participants prospectively from approximately 12-16 weeks of gestation, providing one of the earliest-known cohorts with accurate gestational age determination.[10, 11]

The NFBC1966 dataset consists of 12,231 children born to 12,068 mothers. This cohort represents 96% of all children born in Oulu and Lapland in 1966 with expected delivery dates between Jan 1^st^ and Dec 31^st^ 1966. Blood samples of the children in this cohort were collected for genotyping at age 31 (*i.e.* in 1997) and genetic data was available for 5,402 individuals. Genotyping was completed using Illumina HumanCNV370DUO Analysis BeadChip and the Beadstudio 3.1 algorithm.

The NFBC1986 dataset consists of a prospectively recruited cohort containing 9,432 children born to 9,362 mothers. This cohort represents 99% of all available births in Northern Finland between the 1^st^ of July 1985 and the 30^th^ of June 1986. Blood samples from the children of the NFBC1986 cohort were collected for genotyping at 16 years of age (*i.e.* in 2002-2003). Genetic data is available for 3,739 individuals in total (~500 were selected as representing individuals with GDM, GHT, and preterm birth; the remaining represented a random sample of the cohort). Genotyping was completed using the OmniExpresse Exome Chip and the Beadstudio 3.1 algorithm.

### Defining the post-term dataset

The gestational ages of the individuals in the 1966 and 1986 NFBC cohorts were calculated at their first antenatal visit. For the NFBC1966 cohort gestational age was calculated through last menstrual period. In the NFBC1986 cohort gestational age was based on ultrasound at <20 weeks of gestation or on the last menstral period, with discrepant cases reviewed in detail from medical records as previously described.[44] The control cohort was restricted to those born at full-term which was defined as between 38 0/7 to 40 0/7 weeks of gestation. Children born between 37 0/7 to 37 6/7 weeks (i.e. Early term) or 41 0/7 to 41 6/7 weeks (*i.e.* Late term) were excluded to reduce mischaracterization due to errors in the calculated gestational age. The post-term case cohorts included those individuals born at ≥42 0/7 weeks of gestation. To further reduce the chances of obscuring the genetic potential of gestational age, we excluded individuals from our control cohort who were born early due to induction of labour or other factors: 1) those born from multiple births; 2) those whose mother had gestational diabetes (prediabetes in 1966 cohort); and 3) those whose birth was by planned caesarean section.

### Quality control of genetic data

Genetic data was vetted for quality control. Genetic data for a subject was excluded if: 1) the call rate was < 95% (99 % if the minor allele frequency < 5 %); 2) the mean heterozygosity was < 0.29; 3) there were multidimensional scaling (MDS) outliers; 4) the concordance with other DNA samples in the cohort ≥ 0.99 (risk of being duplicated sample); 5) identity by state (IBS) pairwise comparisons were > 0.99 with most other samples (suspicion of samples being contaminated); 6) IBS pairwise sharing was > 0.20; 7) consent was not given; 8) comparison to medical records identified a gender-genotype mismatch; 9) there was an elevated heterozygosity rate (4 or more standard deviations from the mean); or 10) there was significant deviation from the Hardy-Weinberg Equilibrium (p < 0.0001).

After all QC measures and exclusions were applied, 5,402 and 3,739 individuals remained in the 1966 and 1986 studies, respectively. These included 1034 post-term individuals and 2375 term-born controls from the NFBC1966 cohort and 133 post-term individuals and 1250 term-born controls from the NFBC1986 cohort.

### Imputation of genetic data

Impute version 2 was used to estimate the single nucleotide polymorphisms (SNPs) that were not sampled directly by the genotyping platform for the NFBC 1966 samples. The imputation used HapMap 2 (Build 36) as the reference panel and proper_info > 0.4 as the quality metric[45, 46]. Before imputation, there were 309,948 directly genotyped SNPs. After imputation, 3,855,963 SNPs, including those directly genotyped, were available from the 1966 genotypes for analysis.

Imputation of the missing 1986 genotypes was carried out in two steps: 1) a pre-phasing step that estimated haplotypes for all available samples using the SHAPEIT program with the 1000 genomes reference panel as the guide; and 2) an imputation step (Impute version 2) that imputed the missing alleles directly onto the phased haplotypes[45, 46]. After imputation, 59,683,063 SNPs, including those directly genotyped, were available from the 1986 genotypes for analysis.

### Statistical analysis

SNPTEST version 2[46] was used to perform all genetic analyses on the imputed genetic data for both cohorts. In the regression analysis, the main effects model tested the association between SNP markers and gestational age. SNP genotypes were coded as 0, 1, or 2 (according to the number of copies of the minor allele) and an additive model of genetic variance was assumed where the effect on the trait of the heterozygote was estimated to be midway between the levels of the two homozygotes. This model fits the best assumption for a post-term phenotype that is thought to have a small amount of genetic variance produced by multiple genetic variants in combination.

All genetic analyses also accounted for child’s sex, as this was the only trait that has consistently shown a large effect on post-term birth status in any previous post-term studies[15, 43, 47, 48].

The p-value for results that were suggestive of statistical significance in the discovery phase in each cohort was set at any p-value less than 1 x 10^−5^. In the validation phase, any finding (p < 0.05) within the significant LD block was considered validation of significance of the locus. These p-values were selected because we planned functional analyses to confirm the significant variants,

Quantile-quantile plots were generated (qqman package in R), by plotting the expected distribution of p-values versus the observed p-values (assuming a uniform distribution), to test for possible sources of p-value inflation.

### Spatial analysis of the validated SNPs for putative regulatory roles within the genome

HiC spatial genomic connectivity (HiC) data was used to identify genes that SNPs connected to[24, 27]. GWAS3D[49] was used, with default parameters, to identify physical connections (as captured by proximity ligation) that occurred with the most significant GWAS SNPs.

### Identification of gene expression alterations associated with the GWAS loci

eQTL analysis identifies SNPs that associate with altered expression level(s) of one or more genes[50]. The Genotype-Tissue Expression (GTEx) project database (version 6) eQTL data is powered for the global examination of larger eQTL effects[51]. Therefore, we limited false positives in our trans-eQTL results by only testing eQTLs supported by SNP-gene spatial interactions. Significance levels for this analysis were based on evidence from prior literature[27, 51, 52]: cis-eQTL (genes < 1 Mb distance from SNP, p<1x10^−4^), trans-eQTL (longer distance or inter-chromosomal, p<1x10^−3^). Thus, the identification of a SNP-gene spatial interaction that is re-inforced by SNP-gene eQTL has two independent sources of evidence verifying the long-distance transcription regulatory functions.

### Luciferase assays

Enhancer activity of the post-term SNPS in proximity to loci *(B3GALT5, ARHGAP42, ADAMTS13, SSBP2,* and *TKT)* was measured by luciferase assay. Briefly, the regions spanning each SNP were PCR amplified from genomic DNA obtained from 1000 genome samples cloned into the Gateway adapted pGL4.23-GW (Addgene Plasmid #60323) [53] and sequenced to confirm the genotype. For rs11170213, no sample genotype information could be obtained from the 1000 genomes, so the region spanning the allele ‘A’ of the SNP was amplified from MCF-7 genomic DNA. The Allele ‘C’ version of the SNP was generated by site directed mutageneis (SDM) of the cloned pGL4.23 plasmid using the QuickChange mutageneis protocol (Agilent Technologies). Primers for PCR amplification and SDM are listed in Supplemental Table S1. The *ADAMTS13* locus could not be amplified by PCR. HeLa cells were seeded at 5x10^3^ cells per well in a 96 well plate, grown in DMEM media supplemented with 10% fetal bovine serum (Thermo Scientific, #11995-065) one day prior to transfection. Cells were co-transfected with the cloned pGL4.23 and renilla plasmids using Lipofectamine 3000 (Thermo Scientific, #L3000008) and the the Promega Dual Glo Luciferase Assay System (#PME2920) was used to measure luciferase activity after 48 hours. Luminescence was normalized to Renilla and expressed relative to the normalized luminescence of empty pGL4.23. Results are from four independent biological replicates.

## Acknowledgments

The authors would like to thank and acknowledge the contributions of Tuula Ylitalo, Nikman Adli Nor Hashim, Alexessander Da Silva Couto Alves, Andrianos M Yiorkas, and Minna Männikkö, each of whom played a significant role in data generation and collaboration at the NFBC.

---

**Supplamental Figure S1.**
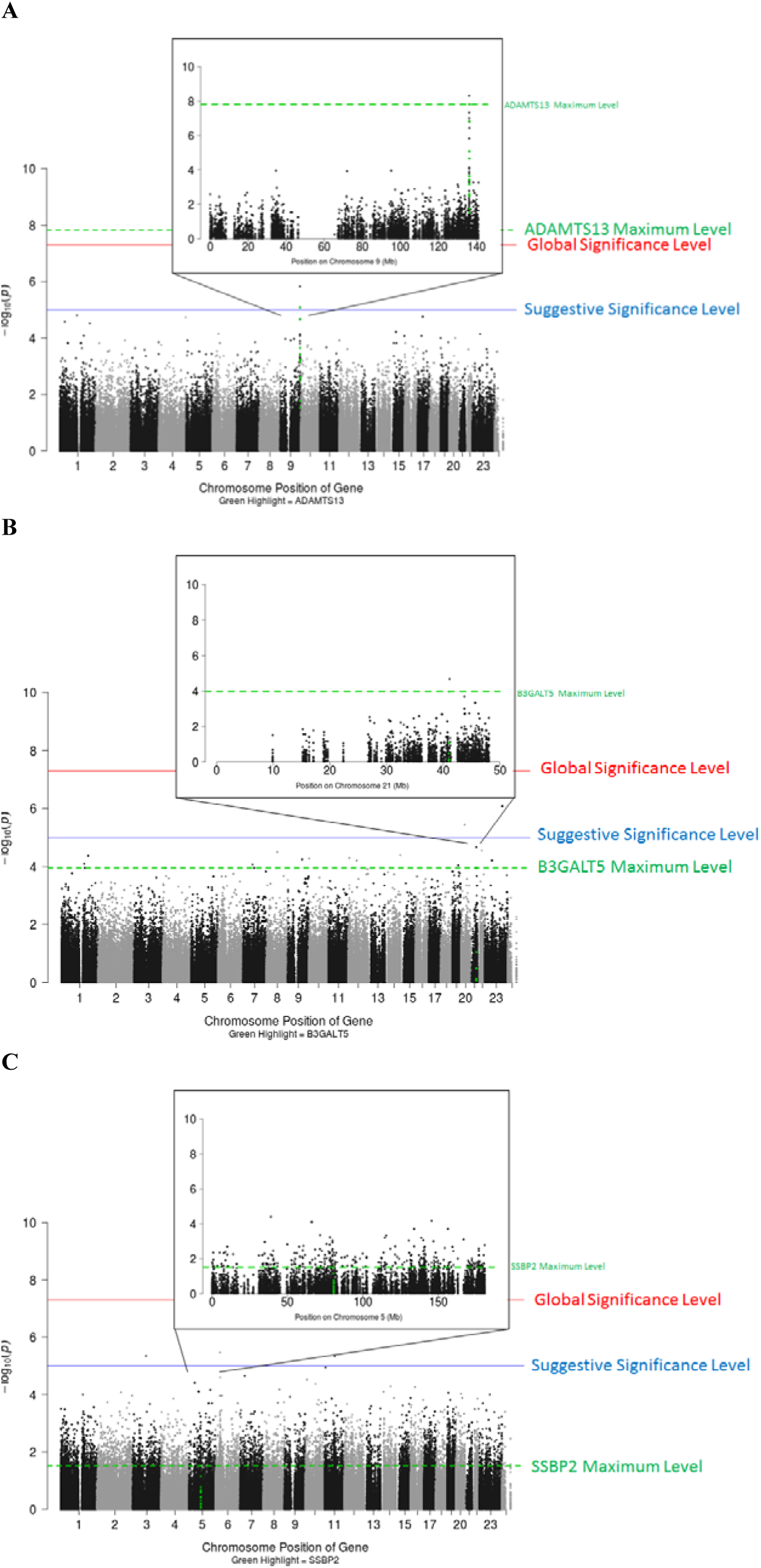
Global eQTL analysis reveals little evidence of trans-eQTLs in the ADAMTS13 (A), B3GALT5 (B), and SSBP2 (C) post-term loci. The −log10 of the eQTL p-values of the association between the ADAMTS13 (A), B3GALT5 (B), and SSBP2 (C) loci and expression in various tissue types in the GTEx database. For rs655911 (ADAMTS13 locus), the only significant peak is at the ADAMTS13 gene (p=1.5x10^−8^). For rs1534080 (B3GALT5 locus), no globally significant (p < 5 x 10^−8^) peaks were identified for, but the highest peak is at the B3GALT5 gene (p=9x10^−5^). For rs2135 (SSBP2 locus), the highest peaks are not at the SSBP2 locus, but are spread out amongst other areas of the genome. Tissue types tested: subcutaneous adipose, aortic artery, tibial artery, heart (left ventricle), lung, tibial nerve, sun-exposed skin (lower leg), skeletal muscle, mucosa and muscularis of the esophagus, thyroid, mammary (breast) tissue, and whole blood.

**Supplemental Table S1.**
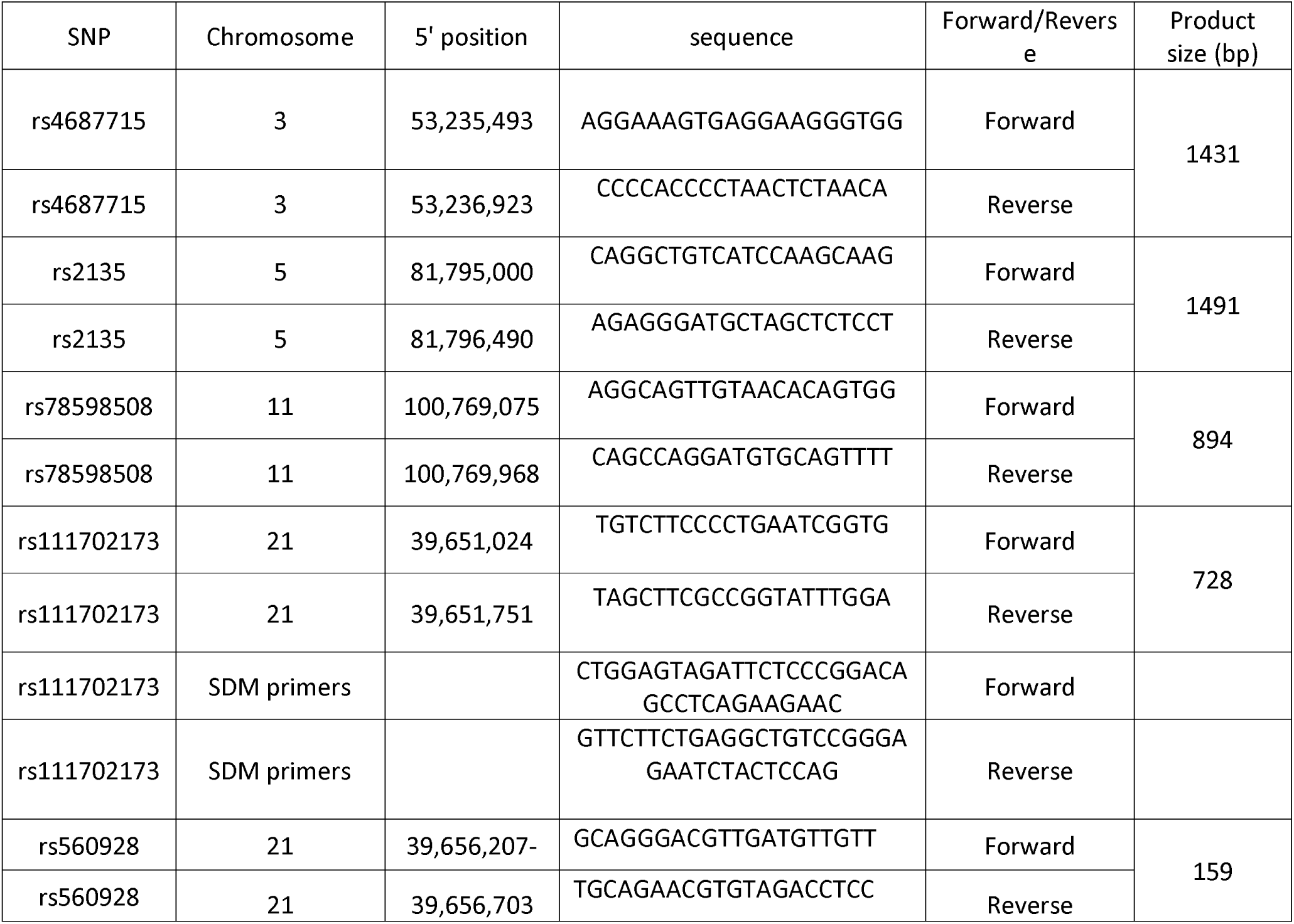
Primer sequences used to amplify genomic DNA regions to test for enhancer activity in the Post-term loci.

## References

1 Duque-Guimaraes DE, Ozanne SE. Nutritional programming of insulin resistance: causes and consequences. Trends Endocrinol Metab 2013;24:525–35.

2 Muglia LI, Katz M. The enigma of spontaneous preterm birth. N Engl J Med 2010;362:529–35.

3 Hofman PL, Regan F, Jackson WE, Jefferies C, Knight DB, Robinson EM, Cutfield WS. Premature birth and later insulin resistance. N Engl J Med 2004;351:2179–86.

4 Sipola-Leppänen M, Vääräsmäki M, Tikanmäki M, Matinolli H-M, Miettola S, Hovi P, Wehkalampi K, Ruokonen A, Sundvall J, Pouta A, Eriksson JG, Järvelin M-R, Kajantie E. Cardiometabolic risk factors in young adults who were born preterm. Am J Epidemiol 2015;181:861–73.

5 Doherty L, Norwitz ER. Prolonged pregnancy: when should we intervene? Curr Opin Obs Gynecol 2008;20:519–27.

6 Ayyavoo A, Derraik JG, Hofman PL, Mathai S, Biggs J, Stone P, Sadler L, Cutfield WS. Prepubertal children born post-term have reduced insulin sensitivity and other markers of the metabolic syndrome. PLoS One 2013;8:e67966.

7 Beltrand J, Soboleva TK, Shorten PR, Derraik JG, Hofman P, Albertsson-Wikland K, Hochberg Z, Cutfield WS. Post-term birth is associated with greater risk of obesity in adolescent males. J Pediatr 2012;160:769–73.

8 Caughey AB, Sundaram V, Kaimal AJ, Cheng YW, Gienger A, Little SE, Lee JF, Wong L, Shaffer BL, Tran SH, Padula A, McDonald KM, Long EF, Owens DK, Bravata DM. Maternal and neonatal outcomes of elective induction of labor. Evid Rep Technol Assess (Full Rep) 2009;:1–257.

9 Järvelin MR, Hartikainen-Sorri AL, Rantakallio P. Labour induction policy in hospitals of different levels of specialisation. Br J Obstet Gynaecol 1993;100:310–5.

10 Pekkanen J, Xu B, Jarvelin MR. Gestational age and occurrence of atopy at age 31--a prospective birth cohort study in Finland. Clin Exp Allergy 2001;31:95–102.

11 Järvelin MR, Elliott P, Kleinschmidt I, Martuzzi M, Grundy C, Hartikainen AL, Rantakallio P. Ecological and individual predictors of birthweight in a northern Finland birth cohort 1986. Paediatr Perinat Epidemiol 1997;11:298–312.

12 Clausson B, Lichtenstein P, Cnattingius S. Genetic influence on birthweight and gestational length determined by studies in offspring of twins. BJOG 2000;107:375–81.

13 Ley GD. Some aspects of prolonged gestation. Med J Aust 1953;2:749–52.

14 Ahn MO, Phelan JP. Epidemiologic aspects of the postdate pregnancy. Clin Obs Gynecol 1989;32:228–34.

15 Kistka ZA, Palomar L, Boslaugh SE, DeBaun MR, DeFranco EA, Muglia LJ. Risk for postterm delivery after previous postterm delivery. Am J Obs Gynecol 2007;196:241 e1-6.

16 Olesen AW, Basso O, Olsen J. Risk of recurrence of prolonged pregnancy. BMJ 2003;326:476.

17 Morken NH, Melve KK, Skjaerven R. Recurrence of prolonged and post-term gestational age across generations: maternal and paternal contribution. BJOG 2011;118:1630–5.

18 Oberg AS, Frisell T, Svensson AC, Iliadou AN. Maternal and fetal genetic contributions to postterm birth: familial clustering in a population-based sample of 475,429 Swedish births. Am J Epidemiol 2013;177:531–7.

19 Laursen M, Bille C, Olesen AW, Hjelmborg J, Skytthe A, Christensen K. Genetic influence on prolonged gestation: a population-based Danish twin study. Am J Obs Gynecol 2004;190:489–94.

20 Schierding W, O’Sullivan JM, Derraik JG, Cutfield WS. Genes and post-term birth: late for delivery. BMC Res Notes 2014;7:720.

21 Welter D, MacArthur J, Morales J, Burdett T, Hall P, Junkins H, Klemm A, Flicek P, Manolio T, Hindorff L, Parkinson H. The NHGRI GWAS Catalog, a curated resource of SNP-trait associations. Nucleic Acids Res 2014;42:D1001–6.

22 Strawbridge RJ, Dupuis J, Prokopenko I, Barker A, Ahlqvist E, Rybin D, Petrie JR, Travers ME, Bouatia-Naji N, Dimas AS, Nica A, Wheeler E, Chen H, Voight BF, Taneera J, Kanoni S, Peden JF, Turrini F, Gustafsson S, Zabena C, Almgren P, Barker DJP, Barnes D, Dennison EM, Eriksson JG, Eriksson P, Eury E, Folkersen L, Fox CS, Frayling TM, Goel A, Gu HF, Horikoshi M, Isomaa B, Jackson AU, Jameson KA, Kajantie E, Kerr-Conte J, Kuulasmaa T, Kuusisto J, Loos RJF, Luan J, Makrilakis K, Manning AK, Mart??nez-Larrad MT, Narisu N, Mannila MN, ??hrvik J, Osmond C, Pascoe L, Payne F, Sayer AA, Sennblad B, Silveira A, Stan??c??kov?? A, Stirrups K, Swift AJ, Syv??nen AC, Tuomi T, Van’t Hlooft FM, Walker M, Weedon MN, Xie W, Zethelius B, Scott LJ, Steinthorsdottir V, Morris AP, Dina C, Welch RP, Zeggini E, Huth C, Aulchenko YS, Thorleifsson G, Mcculloch LJ, Ferreira T, Grallert H, Amin N, Wu G, Willer CJ, Raychaudhuri S, Mccarroll SA, Hofmann OM, Qi L, Segre A V., Van Hoek M, Navarro P, Ardlie K, Balkau B, Benediktsson R, Bennett AJ, Blagieva R, Boerwinkle E, Bonnycastle LL, Bostrom KB, Bravenboer B, Bumpstead S, Burtt NP, Charpentier G, Chines PS, Cornelis M, Couper DJ, Crawford G, Doney ASF, Elliott KS, Elliott AL, Erdos MR, Franklin CS, Ganser M, Gieger C, Grarup N, Green T, Griffin S, Groves CJ, Guiducci C, Hadjadj S, Hassanali N, Herder C, Johnson PR V, Jorgensen T, Kao WHL, Klopp N, Kong A, Kraft P, Lauritzen T, Li M, Lieverse A, Lindgren CM, Lyssenko V, Marre M, Meitinger T, Midthjell K, Morken MA, Nilsson P, Owen KR, Perry JRB, Petersen AK, Platou C, Proenca C, Rathmann W, Rayner NW, Robertson NR, Rocheleau G, Roden M, Sampson MJ, Saxena R, Shields BM, Shrader P, Sigurdsson G, Sparso T, Strassburger K, Stringham HM, Sun Q, Thorand B, Tichet J, Van Dam RM, Van Haeften TW, Van Herpt T, Van Vliet-Ostaptchouk J V., Walters GB, Wijmenga C, Witteman J, Bergman RN, Cauchi S, Collins FS, Gloyn AL, Gyllensten U, Hansen T, Hide WA, Hitman GA, Hofman A, Hunter DJ, Hveem K, Laakso M, Mohlke KL, Morris AD, Palmer CNA, Pramstaller PP, Rudan I, Sijbrands E, Stein LD, Tuomilehto J, Uitterlinden A, Wareham NJ, Watanabe RM, Abecasis GR, Boehm BO, Campbell H, Daly MJ, Hattersley AT, Hu FB, Meigs JB, Pankow JS, Pedersen O, Wichmann HE, Barroso I, Groop L, Sladek R, Thorsteinsdottir U, Wilson JF, Illig T, Froguel P, Van Duijn CM, Stefansson K, Altshuler D, Boehnke M, McCarthy MI, Speliotes EK, Berndt SI, Monda KL, Allen HL, Magi R, Randall JC, Vedantam S, Winkler TW, Workalemahu T, Heid IM, Wood AR, Weyant RJ, Estrada K, Liang L, Nemesh J, Park JH, Kilpelainen TO, Yang J, Esko T, Feitosa MF, Kutalik Z, Mangino M, Scherag A, Smith AV, Welch R, Zhao JH, Aben KK, Absher DM, Dixon AL, Fisher E, Glazer NL, Goddard ME, Heard-Costa NL, Hoesel V, Hottenga JJ, Johansson A, Johnson T, Ketkar S, Lamina C, Li S, Moffatt MF, Myers RH, Peters MJ, Preuss M, Ripatti S, Rivadeneira F, Sandholt C, Timpson NJ, Tyrer JP, Van Wingerden S, White CC, Wiklund F, Barlassina C, Chasman DI, Cooper MN, Jansson JO, Lawrence RW, Pellikka N, Shi J, Thiering E, Alavere H, Alibrandi MTS, Arnold AM, Aspelund T, Atwood LD, Balmforth AJ, Ben-Shlomo Y, Bergmann S, Biebermann H, Blakemore AIF, Boes T, Bornstein SR, Brown MJ, Buchanan TA, Busonero F, Cappuccio FP, Cavalcanti-Proenca C, Chen YDI, Chen CM, Clarke R, Coin L, Connell J, Day INM, Den Heijer M, Duan J, Ebrahim S, Elliott P, Elosua R, Eiriksdottir G, Facheris MF, Felix SB, Fischer-Posovszky P, Folsom AR, Friedrich N, Freimer NB, Fu M, Gaget S, Gejman P V., Geus EJC, Gjesing AP, Goyette P, Grasler J, Greenawalt DM, Gudnason V, Hartikainen AL, Hall AS, Havulinna AS, Hayward C, Heath AC, Hengstenberg C, Hicks AA, Hinney A, Homuth G, Hui J, Igl W, Iribarren C, Jacobs KB, Jarick I, Jewell E, John U, Jousilahti P, Jula A, Kaakinen M, Kaplan LM, Kathiresan S, Kettunen J, Kinnunen L, Knowles JW, Kolcic I, K??nig IR, Koskinen S, Kovacs P, Kvaloy K, Laitinen J, Lantieri O, Lanzani C, Launer LJ, Lecoeur C, Lehtimaki T, Lettre G, Liu J, Lokki ML, Lorentzon M, Luben RN, Ludwig B, Manunta P, Marek D, Martin NG, McArdle WL, McCarthy A, McKnight B, Melander O, Meyre D, Montgomery GW, Mulic R, Ngwa JS, Nelis M, Neville MJ, Nyholt DR, O’Donnell CJ, O’Rahilly S, Ong KK, Oostra B, Pare G, Parker AN, Perola M, Pichler I, Pietilainen KH, Platou CGP, Polasek O, Pouta A, Rafelt S, Raitakari O, Rayner NW, Ridderstrale M, Rief W, Ruokonen A, Rzehak P, Salomaa V, Sanders AR, Sandhu MS, Sanna S, Saramies J, Savolainen MJ, Scherag S, Schipf S, Schreiber S, Schunkert H, Silander K, Sinisalo J, Siscovick DS, Smit JH, Soranzo N, Sovio U, Stephens J, Surakka I, Tammesoo ML, Tardif JC, Teder-Laving M, Teslovich TM, Thompson JR, Thomson B, Tonjes A, Van Meurs JBJ, Van Ommen GJ, Vatin V, Viikari J, Visvikis-Siest S, Vitart V, Vogel CIG, Waite LL, Wallaschofski H, Widen E, Wiegand S, Wild SH, Willemsen G, Witte DR, Witteman JC, Xu J, Zhang Q, Zgaga L, Ziegler A, Zitting P, Beilby JP, Farooqi IS, Hebebrand J, Huikuri H V., James AL, Kahonen M, Levinson DF, Macciardi F, Nieminen MS, Ohlsson C, Palmer LJ, Ridker PM, Stumvoll M, Beckmann JS, Boeing H, Boomsma DI, Caulfield MJ, Chanock SJ, Cupples LA, Smith GD, Erdmann J, Gronberg H, Hall P, Harris TB, Hayes RB, Heinrich J, Jarvelin MR, Kaprio J, Karpe F, Khaw KT, Kiemeney LA, Krude H, Lawlor DA, Metspalu A, Munroe PB, Ouwehand WH, Penninx BW, Peters A, Quertermous T, Reinehr T, Rissanen A, Samani NJ, Schwarz PEH, Shuldiner AR, Spector TD, Uda M, Valle TT, Wabitsch M, Waeber G, Watkins H, Wright AF, Zillikens MC, Chatterjee N, McCarroll SA, Purcell S, Schadt EE, Visscher PM, Assimes TL, Borecki IB, Deloukas P, Groop LC, Haritunians T, Kaplan RC, O’Connell JR, Peltonen L, Schlessinger D, Strachan DP, Wichmann HE, North KE, Hirschhorn JN, Ingelsson E, Nica AC, Parts L, Glass D, Nisbet J, Barrett A, Sekowska M, Travers M, Potter S, Grundberg E, Small K, Hedman AK, Bataille V, Bell JT, Surdulescu G, Ingle C, Nestle FO, Di Meglio P, Min JL, Wilk A, Hammond CJ, Yang TP, Montgomery SB, O’Rahilly S, Zondervan KT, Durbin R, Ahmadi K, Dermitzakis ET, Reilly MP, Holm H, Stewart AFR, Barbalic M, Absher D, Aherrahrou Z, Allayee H, Anand SS, Andersen K, Anderson JL, Ardissino D, Ball SG, Barnes TA, Becker DM, Becker LC, Berger K, Bis JC, Boekholdt SM, Braund PS, Burnett MS, Buysschaert I, Carlquist JF, Chen L, Cichon S, Codd V, Davies RW, Dedoussis G, Dehghan A, Demissie S, Devaney JM, Diemert P, Do R, Doering A, Eifert S, El Mokhtari NE, Ellis SG, Engert JC, Epstein SE, De Faire U, Fischer M, Freyer J, Gigante B, Girelli D, Gretarsdottir S, Gulcher JR, Halperin E, Hammond N, Hazen SL, Horne BD, Jones GT, Jukema JW, Kaiser MA, Kastelein JJP, Kolovou G, Laaksonen R, Lambrechts D, Leander K, Li M, Lieb W, Loley C, Lotery AJ, Mannucci PM, Maouche S, Martinelli N, McKeown PP, Meisinger C, Merlini PA, Mooser V, Morgan T, M??hleisen TW, Muhlestein JB, M??nzel T, Musunuru K, Nahrstaedt J, Nelson CP, N??then MM, Olivieri O, Patel RS, Patterson CC, Peyvandi F, Qu L, Quyyumi AA, Rader DJ, Rallidis LS, Rice C, Rosendaal FR, Rubin D, Sampietro ML, Sandhu MS, Schadt E, Sch??fer A, Schillert A, Schrezenmeir J, Schwartz SM, Sivananthan M, Sivapalaratnam S, Smith A, Smith TB, Snoep JD, Spertus JA, Stark K, Stoll M, Wilson Tang WH, Tennstedt S, Thorgeirsson G, Tomaszewski M, Uitterlinden AG, Van Rij AM, Wareham NJ, Wells GA, Wild PS, Willenborg C, Witteman JCM, Wright BJ, Ye S, Zeller T, Cambien F, Goodall AH, Marz W, Blankenberg S, Roberts R, McPherson R, Hopewell JC, Parish S, Offer A, Bowman L, Sleight P, Armitage J, Peto R, Collins R, Chambers JC, Abecasis G, Ahmed N, Caulfield M, Donnelly P, Kooner AS, McCarthy M, Samani N, Scott J, Sehmi J, Zhang W, Kooner JS, Strawbridge R, Sabater-Lleal M, M??larstig A, Hell??nius ML, Van’t Hooft F, Olsson G, Rust S, Assmann G, Seedorf U, Barlera S, Tognoni G, Franzosi MG, Linksted P, Ongen H, Kyriakou T, Green F, Farrall M, Saleheen D, Rasheed A, Zaidi M, Shah N, Samuel M, Mallick N, Azhar M, Zaman K, Samad A, Ishaq M, Gardezi A, Memon FUR, Frossard P, Danesh J, Chambers J, Kooner J, ??stenson CG, Lind L, Cooper CC, Serrano-R??os M, Ferrannini E, Forsen TJ, Johnson P, Pattou F, Dedoussis G V., Langenberg C, Hamsten A, Florez JC. Genome-wide association identifies nine common variants associated with fasting proinsulin levels and provides new insights into the pathophysiology of type 2 diabetes. Diabetes 2011;60:2624–34.

23 Schierding W, Cutfield WS, O’Sullivan JM. The missing story behind Genome Wide Association Studies: single nucleotide polymorphisms in gene deserts have a story to tell. Front Genet 2014;5:39.

24 Schierding W, O’Sullivan JM. Connecting SNPs in diabetes: A spatial analysis of meta-GWAS loci. Front Endocrinol (Lausanne) 2015;6. doi:10.3389/fendo.2015.00102

25 Smemo S, Tena JJ, Kim K-H, Gamazon ER, Sakabe NJ, Gómez-Marín C, Aneas I, Credidio FL, Sobreira DR, Wasserman NF, Lee JH, Puviindran V, Tam D, Shen M, Son JE, Vakili NA, Sung HK, Naranjo S, Acemel RD, Manzanares M, Nagy A, Cox NJ, Hui C-C, Gomez-Skarmeta JL, Nóbrega MA. Obesity-associated variants within FTO form long-range functional connections with IRX3. Nature 2014;507:371–5.

26 Kelemen LE, Lawrenson K, Tyrer J, Li Q, Lee JM, Seo J-H, Phelan CM, Beesley J, Chen X, Spindler TJ, Aben KKH, Anton-Culver H, Antonenkova N. Genome-wide significant risk associations for mucinous ovarian carcinoma. Nat Genet 2015;47:888–97.

27 Schierding W, Antony J, Cutfield WS, Horsfield JA, O’Sullivan JM. Intergenic GWAS SNPs are key components of the spatial and regulatory network for human growth. Hum Mol Genet 2016;25:3372–82.

28 Rey E, Kahn SR, David M, Shrier I. Thrombophilic disorders and fetal loss: a meta-analysis. Lancet (London, England) 2003;361:901–8.

29 McMillen IC, Robinson JS. Developmental origins of the metabolic syndrome: prediction, plasticity, and programming. Physiol Rev 2005;85:571–633.

30 Kim Y, Kim E-Y, Seo Y-M, Yoon TK, Lee W-S, Lee K-A. Function of the pentose phosphate pathway and its key enzyme, transketolase, in the regulation of the meiotic cell cycle in oocytes. Clin Exp Reprod Med 2012;39:58–67.

31 Kaya S, Keskin HL, Kaya B, Ustuner I, Avsar AF. Reduced total antioxidant status in postterm pregnancies. Hippokratia 2013;17:55–9.

32 Xu Z-P, Wawrousek EF, Piatigorsky J. Transketolase Haploinsufficiency Reduces Adipose Tissue and Female Fertility in Mice. Mol Cell Biol 2002;22:6142–7.

33 Karotia HC, Inamdar S, Khan MA, Mathur PS. Foetal haemoglobin concentration in cord blood in relation to gestational age. Indian J Pediatr 1976;43:313–8.

34 Glasser L, Sutton N, Schmeling M, Machan JT. A comprehensive study of umbilical cord blood cell developmental changes and reference ranges by gestation, gender and mode of delivery. J Perinatol 2015;35:469–75.

35 Tuckuviene R, Christensen AL, Helgested J, Hundborg HH, Kristensen SR, Johnsen SP. Infant, obstetrical and maternal characteristics associated with thromboembolism in infancy: a nationwide population-based case-control study. Arch Dis Child Fetal Neonatal Ed 2012;97:F417-422.

36 Groudine M, Kohwi-Shigematsu T, Gelinas R, Stamatoyannopoulos G, Papayannopoulou T. Human fetal to adult hemoglobin switching: changes in chromatin structure of the betaglobin gene locus. Proc Natl Acad Sci U S A 1983;80:7551–5.

37 Astorga GP, Williams RC. Altered reactivity in mixed lymphocyte culture of lymphocytes from patients with rheumatoid arthritis. Arthritis Rheum 1969;12:547–54.

38 Hamamy H, Makrythanasis P, Al-Allawi N, Muhsin AA, Antonarakis SE. Recessive thrombocytopenia likely due to a homozygous pathogenic variant in the FYB gene: case report. BMC Med Genet 2014;15:135.

39 Peterson EJ, Woods ML, Dmowski SA, Derimanov G, Jordan MS, Wu JN, Myung PS, Liu QH, Pribila JT, Freedman BD, Shimizu Y, Koretzky GA. Coupling of the TCR to integrin activation by Slap-130/Fyb. Science 2001;293:2263–5.

40 Majerus EM, Anderson PJ, Sadler JE. Binding of ADAMTS13 to von Willebrand factor. J Biol Chem 2005;280:21773–8.

41 Ricketts LM, Dlugosz M, Luther KB, Haltiwanger RS, Majerus EM. O-Fucosylation Is Required for ADAMTS13 Secretion. J Biol Chem 2007;282:17014–23.

42 NICE. Induction of labour clinical guideline 70. 2008;2013.

43 Myklestad K, Vatten LJ, Magnussen EB, Salvesen KA, Romundstad PR. Do parental heights influence pregnancy length?: a population-based prospective study, HUNT 2. BMC Pregnancy Childbirth 2013;13:33.

44 Sipola-Leppanen M, Vaarasmaki M, Tikanmaki M, Hovi P, Miettola S, Ruokonen A, Pouta A, Jarvelin M-R, Kajantie E. Cardiovascular Risk Factors in Adolescents Born Preterm. Pediatrics 2014;134:e1072–81.

45 Howie BN, Donnelly P, Marchini J. A flexible and accurate genotype imputation method for the next generation of genome-wide association studies. PLoS Genet 2009;5:e1000529.

46 Marchini J, Howie B. Genotype imputation for genome-wide association studies. Nat Rev Genet 2010;11:499–511.

47 Stotland NE, Washington AE, Caughey AB. Prepregnancy body mass index and the length of gestation at term. Am J Obs Gynecol 2007;197:378 e1-5.

48 Caughey AB, Stotland NE, Washington AE, Escobar GJ. Who is at risk for prolonged and postterm pregnancy? Am J Obs Gynecol 2009;200:683 e1-5.

49 Li MJ, Wang LY, Xia Z, Sham PC, Wang J. GWAS3D: Detecting human regulatory variants by integrative analysis of genome-wide associations, chromosome interactions and histone modifications. Nucleic Acids Res 2013;41:W150-8.

50 Albert FW, Kruglyak L. The role of regulatory variation in complex traits and disease. Nat Rev Genet 2015;16:197–212.

51 Ardlie KG, Deluca DS, Segre A V., Sullivan TJ, Young TR, Gelfand ET, Trowbridge CA, Maller JB, Tukiainen T, Lek M, Ward LD, Kheradpour P, Iriarte B, Meng Y, Palmer CD, Esko T, Winckler W, Hirschhorn JN, Kellis M, MacArthur DG, Getz G, Shabalin AA, Li G, Zhou Y-H, Nobel AB, Rusyn I, Wright FA, Lappalainen T, Ferreira PG, Ongen H, Rivas MA, Battle A, Mostafavi S, Monlong J, Sammeth M, Mele M, Reverter F, Goldmann JM, Koller D, Guigo R, McCarthy MI, Dermitzakis ET, Gamazon ER, Im HK, Konkashbaev A, Nicolae DL, Cox NJ, Flutre T, Wen X, Stephens M, Pritchard JK, Tu Z, Zhang B, Huang T, Long Q, Lin L, Yang J, Zhu J, Liu J, Brown A, Mestichelli B, Tidwell D, Lo E, Salvatore M, Shad S, Thomas JA, Lonsdale JT, Moser MT, Gillard BM, Karasik E, Ramsey K, Choi C, Foster BA, Syron J, Fleming J, Magazine H, Hasz R, Walters GD, Bridge JP, Miklos M, Sullivan S, Barker LK, Traino HM, Mosavel M, Siminoff LA, Valley DR, Rohrer DC, Jewell SD, Branton PA, Sobin LH, Barcus M, Qi L, McLean J, Hariharan P, Um KS, Wu S, Tabor D, Shive C, Smith AM, Buia SA, Undale AH, Robinson KL, Roche N, Valentino KM, Britton A, Burges R, Bradbury D, Hambright KW, Seleski J, Korzeniewski GE, Erickson K, Marcus Y, Tejada J, Taherian M, Lu C, Basile M, Mash DC, Volpi S, Struewing JP, Temple GF, Boyer J, Colantuoni D, Little R, Koester S, Carithers LJ, Moore HM, Guan P, Compton C, Sawyer SJ, Demchok JP, Vaught JB, Rabiner CA, Lockhart NC. The Genotype-Tissue Expression (GTEx) pilot analysis: Multitissue gene regulation in humans. Science 2015;348:648–60.

52 Westra H-J, Peters MJ, Esko T, Yaghootkar H, Schurmann C, Kettunen J, Christiansen MW, Fairfax BP, Schramm K, Powell JE, Zhernakova A, Zhernakova D V, Veldink JH, Van den Berg LH, Karjalainen J, Withoff S, Uitterlinden AG, Hofman A, Rivadeneira F, ’t Hoen PAC, Reinmaa E, Fischer K, Nelis M, Milani L, Melzer D, Ferrucci L, Singleton AB, Hernandez DG, Nalls MA, Homuth G, Nauck M, Radke D, Völker U, Perola M, Salomaa V, Brody J, Suchy-Dicey A, Gharib SA, Enquobahrie DA, Lumley T, Montgomery GW, Makino S, Prokisch H, Herder C, Roden M, Grallert H, Meitinger T, Strauch K, Li Y, Jansen RC, Visscher PM, Knight JC, Psaty BM, Ripatti S, Teumer A, Frayling TM, Metspalu A, van Meurs JBJ, Franke L. Systematic identification of trans eQTLs as putative drivers of known disease associations. Nat Genet 2013;45:1238–43.

53 Pasquali L, Gaulton KJ, Rodriguez-Segui SA, Mularoni L, Miguel-Escalada I, Akerman I, Tena JJ, Moran I, Gomez-Marin C, van de Bunt M, Ponsa-Cobas J, Castro N, Nammo T, Cebola I, Garcia-Hurtado J, Maestro MA, Pattou F, Piemonti L, Berney T, Gloyn AL, Ravassard P, Gomez-Skarmeta JL, Muller F, McCarthy MI, Ferrer J, Rodríguez-Seguí SA, Mularoni L, Miguel-Escalada I, Akerman I, Tena JJ, Morán I, Gómez-Marín C, van de Bunt M, Ponsa-Cobas J, Castro N, Nammo T, Cebóla I, García-Hurtado J, Maestro MA, Pattou F, Piemonti L, Berney T, Gloyn AL, Ravassard P, Gómez-Skarmeta JL, Müller F, McCarthy MI, Ferrer J. Pancreatic islet enhancer clusters enriched in type 2 diabetes risk-associated variants. Not Genet 2014;46:136–43.

